# CST–Polα/primase, backed up by Rif1, protects telomeric ss-dsDNA junctions from damage recognition and resection

**DOI:** 10.64898/2026.10.22.760787

**Authors:** Eun Young Yu, Kimberly Calugaru, Tenzin Younten, Sophie Huang, Neal F. Lue

## Abstract

The yeast Cdc13-Stn1-Ten1 (CST) complex is critical for telomere length regulation and telomere protection. Recent studies further implicate Polα/primase (PP) as an essential partner of CST in executing these functions, but the mechanistic bases remain poorly understood. Using the pathogenic yeast *Nakaseomyces glabratus*, a model amenable to mechanistic dissection of CST-PP, we analyzed the protection mechanisms of CST-PP by combining mutations that disrupt individual subunits or complex assembly with mutations in DNA damage response (DDR) and DNA repair factors. The complete loss of Cdc13 – which eliminates the CST-PP assembly – results in telomere damage recognition involving Rad9, Rad24 and Mec3, as well as aberrant resection mediated by Exo1 and Sgs1. In contrast, disrupting the interaction between CST and PP leads to proliferation defects triggered primarily by Exo1, implying a more selective protection function for PP. The involvement of the DDR clamp loader and clamp together with Exo1 points to ss-dsDNA junctions with recessed 5’ ends as a crucial lesion in the absence of Cdc13. Further analysis implicates RNase H1 and H2 in promoting DNA damage, possibly by exposing such junctions. Based on similarities between Cdc13 and POT1, a telomere protein previously implicated in junctional protection, we modeled Cdc13 binding to a ss-dsDNA junction and showed that point mutations at the proposed interaction site impaired telomere protection and altered the distribution of the telomere C-strand 5’ ends *in vivo*. We further demonstrate that Rif1, another telomere protein with junctional binding activity, supports the protection function of CST-PP. Together, our findings illuminate the molecular basis of telomere DNA damage and of CST-PP mediated telomere protection.

## Introduction

Telomeres, the specialized nucleoprotein structures at the tips of eukaryotic chromosomes, promote genome integrity by protecting chromosome ends against aberrant rearrangements (1,2). The telomere nucleoprotein assembly consists of many copies of a short, repeated DNA sequence – with a G-rich 3’ strand and a C-rich 5’ strand – bound by proteins recruited via DNA-protein and protein-protein interactions (3). While most telomere DNA is double-stranded (ds), the most distal region comprises a single-stranded (ss) 3’ overhang (G-tail) that is also covered by protective proteins. Aside from end protection, telomeres rely on specialized DNA synthesis complexes to minimize DNA loss during replication and to add DNA repeats onto chromosome ends, thereby overcoming the end replication problem (4–6). Notably, telomere DNA maintenance is itself strictly regulated, because excessive elongation can also compromise telomere function.

The protection and DNA maintenance functions of telomeres are somewhat at odds: the former shields the telomere terminus from the DNA repair machineries, whereas the latter generates intermediates – ssDNA and ss-dsDNA junctions – that those same machineries recognize. One solution to this dilemma is to use dual-function proteins that promote DNA synthesis while protecting the DNA they generate. Emerging evidence suggests that Cdc13-Stn1-Ten1-Polα/primase (CST-PP), a binary assembly of a single-stranded DNA-binding complex (CST) and a polymerase (PP), exemplifies this solution (7–9). First, CST-PP performs C-strand fill-in synthesis on the lagging strand following disassembly of the replication fork, minimizing DNA loss. Second, in the two-step telomere lengthening pathway, CST-PP synthesizes the telomere C-strand after telomerase has elongated the G-strand, and in doing so also restricts G-strand elongation by displacing telomerase (10,11). Third, CST-PP serves essential protection functions by suppressing the DNA damage response and aberrant repair at telomeres (8,12,13).

That CST-PP executes these functions is supported by substantial biochemical and genetic evidence. *In vitro*, CST binds the telomere G-strand with high affinity and sequence specificity, consistent with CST’s structural similarities to RPA, the ubiquitous ssDNA-binding complex in eukaryotes (10,14). CST also stimulates PP polymerization on both telomeric and non-telomeric substrates, and CST alone can inhibit telomerase by sequestering the G-strand primer (10,11,15,16). Mammalian CST inhibits EXO1 and BLM/DNA2, the two major long-range resection activities at double-strand breaks (DSBs) (17). *In vivo*, mutations in CST and PP subunits cause telomere shortening or lengthening in a mutation- and context-dependent manner (7,9). Yeast CST evidently provides a critical telomere protection function as well: all three subunits are essential, and *ts* alleles of each gene induce a variety of telomere damage and cell cycle defects at semi-permissive and non-permissive temperatures (18–20). Notably, this protection function appears to be replication-dependent; telomere degradation and instability occur only when Cdc13-deficient cells progress through S-phase (12,21). A protection function for PP is less clear and more challenging to demonstrate, given PP’s essential role in chromosomal replication, but recent studies indicate that disrupting PP’s interaction with CST can induce abnormal telomere resection (8). Mammalian PP mutations also elevate ssDNA levels, suggesting loss of protection (22).

Despite the accumulated findings, central questions remain. First, although multiple DDR and repair pathways are evidently activated in the absence of CST-PP, the lesions that activate them – and the reason these lesions arise only during replication – have not been delineated. Second, the mechanisms by which individual CST-PP subunits and the full complex antagonize aberrant repair remain elusive. Third, the mechanistic relationship between the telomere length regulation and telomere protection functions of CST-PP is obscure; the CST *ts* mutants consistently display elongation defects at lower temperatures and protection defects at higher ones, which suggests coupling, yet no molecular model accounts for it. Finally, many other proteins and complexes are implicated in telomere length regulation and protection, and how CST-PP cooperates with them is incompletely understood (23,24). Yeast Rif1 presents a particularly interesting mechanistic and evolutionary puzzle in this regard. Mammalian RIF1 is an upstream regulator of CST-PP at DSBs but has not been implicated in telomere length regulation (25,26), whereas yeast Rif1 regulates DSB repair, telomere length and telomere protection yet does not appear to interact directly with CST-PP (27–30).

We have developed *Nakaseomyces glabratus* (previously named *Candida glabrata*) as a tractable model system for investigating CST-PP mechanisms *in vitro* and *in vivo*. We established co-expression/purification systems for *N. glabratus* (*Ng*) CST and PP as well as *in vitro* assays for their biochemical activities (31), and showed that CST stimulates PP by enhancing both the priming and the primase-to-polymerase switch reactions (15). More recently, by leveraging cryo-EM structures of human CST-PP and AlphaFold modeling, we identified separation-of-function mutations that specifically disrupt the formation of a CST-PP-DNA complex, designated the pre-initiation complex (PIC). These disruptions abolished the stimulatory effect of CST on PP activity *in vitro* and induced abnormal telomere elongation or loss of protection *in vivo*, reinforcing the utility of the *N. glabratus* system and implicating PP as a critical partner of CST. Herein we extend this work to the protection mechanism of *Ng*CST-PP. Using a system that conditionally eliminates Cdc13, we show that its absence triggers telomere DNA damage recognition involving at least Rad9, Rad24 and Mec3, as well as aberrant resection mediated by both Exo1 and Sgs1. In contrast, selectively disrupting the PIC causes damage mediated primarily by Exo1, with much less contribution from Sgs1. These results reveal a division of labor within CST-PP with respect to telomere protection. In addition, the involvement of Rad24, Mec3 and Exo1 in Cdc13-deficient cells points to the exposure of ss-dsDNA junctions (SDJ) as the underlying lesion. We further implicate RNase H1 and H2 in exposing such junctions through degradation of the RNA primers of Okazaki fragments. Guided by the structural and functional similarities between Cdc13 and POT1 (32), we identified point mutations in Cdc13 that impair telomere protection and alter the telomere C-strand 5’ ends. Finally, we show that Rif1, another telomere protein with junctional binding activity, supports the protection function of CST-PP.

## Methods

### *Nakaseomyces glabratus* strain construction and growth

Standard protocols were employed for the genetic manipulation of *N. glabratus* (previously *Candida glabrata*). All strains used in this study are derived from the BG14 strain background (33) and are listed in Supp. Table 1.

The construction of the *exo1Δ* and *rad24Δ* mutants was previously described (8). The *cdc13Δ*, *mec3Δ, rad9Δ, rif1Δ* and *sgs1Δ* mutations were introduced by integrating an NAT^R^ cassette flanked by ∼600 – 700 bp of 5’ UTR and 3’ UTR derived from the corresponding genes. The UTR fragments were generated by PCR amplification and inserted, by either restriction digestion and ligation or Gibson Assembly, between the *Sac*II and *Hpa*I sites and between the *Aat*II and *Pvu*II sites of pBV65 (34) (see Supp. Table 2 for the oligonucleotides used). The cassettes were then released from the plasmid by *Aat*II and *Hpa*I cleavage and used in transformation. Correct clones were selected on media containing 100 µg/ml nourseothricin and identified by PCR genotyping. The NAT^R^ marker was then excised by growing the strains in SD-met supplemented with 1 µM estradiol overnight to induce Cre expression. Cultures were plated on non-selective media (i.e., without nourseothricin) and individual colonies were tested for loss of the NAT^R^ marker.

The pCU-Cdc13-FG_3_ and pCN-Cdc13-FG_3_ plasmids were constructed by first inserting the *Ng*Cdc13 ORF in frame and upstream of the FG_3_ tag of pSMT3-FG_3_ plasmid (pSMT3-Cdc13-FG_3_), and then placing the Cdc13-FG_3_ fragment in between the XmaI and XhoI sites of pCU-PDC1 and pCN-PDC1. Mutant alleles of Cdc13 (Cdc13^R411E^, Cdc13^R531E^ and Cdc13^RREE^) were generated by PCR mutagenesis using pSMT3-Cdc13-FG_3_ and then used to replace the WT ORF in pCU-Cdc13-FG_3_ and pCN-Cdc13-FG_3_. For plasmid eviction, *cdc13Δ* strains harboring pCU-Cdc13-FG_3_ (designated *cdc13Δ\**) were inoculated into YPD and grown overnight, followed by plating on FOA plates or growth in FOA liquid medium. For plasmid shuffle, *cdc13Δ\** was first transformed with pCN-Cdc13-FG_3_ carrying wild type or mutant alleles, passaged in YPD liquid media, then streaked on FOA plates to select for the loss of pCU-Cdc13-FG_3_.

To generate a point mutant of Cdc13 at the native locus, the mutant allele (fused to a C-terminal FLAG_3_ tag) was cloned between a *Not*I site (at nucleotide position 6) and the *Pvu*II site of pBV65, and a *CDC13* 3’ UTR fragment was cloned between the second *Not*I site (at nucleotide position 5712) and its neighboring LoxP site. The integration cassette was released with *Not*I cleavage and used to transform the conditional *cdc13Δ* strain. Correctly reconstituted strains were subjected to NAT^R^ marker recycling, grown in YPD, and then selected on FOA media for the loss of pCU-Cdc13-FG_3_.

For growth analysis, liquid cultures of *Ng* strains were first diluted to OD_600_ of 2.0 (corresponding to ∼ 6 x 10^7^ cells/ml) and subjected to serial 6-fold dilutions. Four microliters of each dilution were spotted onto semi-solid media and grown at 30 °C for up to 3 days, with successive image acquisitions.

### Telomere DNA analyses

Southern analysis of telomere restriction fragments (TRF Southern) was performed using DNA treated with *Eco*RI or *Eco*RI plus *Alu*I as described previously (8,35). The blots were hybridized to a 5’ ^32^P-labeled oligonucleotide containing three copies of the *C. glabrata* C-strand repeat (CgTEL-C3; [GCACCCAGACCCCACA]_3_) and then analyzed by PhosphorImager scanning. Alternatively, the blots were hybridized to a biotinylated G or C-strand probe followed by detection with Streptavidin conjugated-horse radish peroxidase and chemiluminescence. Briefly, the biotinylated probes were synthesized according to (36) and (37) except that (1) DIG-dUTP was replaced by Biotin-11-dUTP; and (2) the human telomere G- and C-strand sequences in the templates were replaced by three copies of the *C. glabrata* telomere repeats (see Supp. Table 2 for the template and primer oligonucleotides). Following hybridization and washing, the blots were incubated in blocking buffer (1x MAB [100 mM maleic acid, 150 mM NaCl, pH 7.5], 1% casein) at room temperature for 1 h with gentle agitation, followed by incubation in streptavidin-HRP Buffer (1x MAB, 0.5% casein, 0.3% Tween-20, 1: 10,000 dilution of HRP-Streptavidin [GenScript, Inc.]) for 1 h. The blots were then washed three times in HRP-SA washing solution (1× MAB, 0.3% Tween-20) and twice in HRP-SA High salt washing solution (1× MAB, 0.5M NaCl, 0.3% Tween-20), followed by exposure to chemiluminescent substrates (Tanon Life Science Co.) and image collection on ChemiDoc MP. C-circle analysis was performed as before with minor modifications (35). Genomic DNAs (12 ng, digested with *Eco*RI) were added to 20 µl mixture containing 4 µg/ml BSA, 4 mM DTT, 0.1% Tween, 1 mM each of dCTP, dGTP and dTTP, 1× NxGen Φ29 Buffer and 5 U NxGen Φ29 DNA polymerase (LGC Inc.) and incubated at 30 °C for 16 h. The DNA polymerase was then inactivated at 65 °C for 20 min. The reaction products were dot-blotted onto a 2x SSC-soaked nylon membrane and the DNAs were UV-cross-linked onto the membrane. The blots were hybridized at 63 °C with ^32^P-labeled CgTEL-C3, and exposed to Phosphor plates. Signals were collected using a PhosphorImager (Amersham Typhoon IP) and quantified with ImageQuant software.

For mapping the C-strand 5’ ends, the STELA protocols previously used to characterize individual telomeres in human and *U. maydis* cells were adapted to the *N. glabratus* system (38,39). Briefly, 5 ng of *Eco*RI-digested genomic DNAs were ligated to individual telorette primers (0.0001 µM) in 15 µl reaction containing 40 U/µl T4 DNA ligase at 35°C for 20 hours. One tenth of the reaction products were then used as templates in 15 µl PCR reactions containing 1x Failsafe PCR Mix H, 1 µM each of the subtelomeric forward primer and the Teltail reverse primer, and 0.75 U Failsafe polymerase. The thermocycling were performed for either 26 or 31 cycles using the following parameters: 30 sec at 94°C, 30 sec at 63°C and 2 min at 72°C. The 26-cycle reaction products were quantified by Southern and PhosphorImager and the 31-cycle reaction products were quantified by gel electrophoresis and ethidium bromide staining, respectively.

### Co-immunoprecipitation (co-IP) and Chromatin immunoprecipitation (ChIP) analysis

For Co-IP, *N. glabratus* strains carrying chromosomally tagged Rif1-Myc_5_ and/or Cdc13-FG_3_ were cultured to OD_600_ of 5.0 and extracts were prepared by lysing cells with glass beads in lysis buffer (50 mM Tris, pH 7.5, 150 mM NaCl, 1 % Triton X-100, 0.1% sodium deoxycholate). Two milligrams of each extract in 500 µl of lysis buffer (with 50 µg of BSA and additional NaCl to a final concentration of 300 mM) was subjected to IP using 15 µl of Anti-DYKDDDDK G1 Affinity Resin beads (GenScript) at 4 °C overnight. IP samples were washed three times at 4 °C for 5 min with rotation in washing buffer (50 mM Tris, pH 7.5, 500 mM NaCl, 10 % glycerol, 2.5 mM MgCl_2_, 1 mM DTT, 0.1% NP-40). All lysis and wash buffers contained protease inhibitors. Bead-bound proteins were eluted by boiling in SDS loading buffer and analyzed by western blotting using Myc antibodies (9E10) (Covance) and FLAG antibodies (GenScript).

For ChIP analysis, cells were crosslinked at OD_600_ of 1.0 with 1 % formaldehyde for 30 min at 30°C, and the reaction was quenched with 125 mM glycine for 5 min at 30 °C. Formaldehyde-fixed or untreated cells were resuspended in lysis buffer (50 mM Tris, pH 7.5, 300 mM NaCl,1 % Triton X-100, 0.1 % sodium deoxycholate) and broken by glass beads. The lysates were sonicated 36 times for 5 s each (constant duty cycle, 40% output) to shear DNAs to a mean length of ∼600 bp. Two milligrams of each extract in 500 µl of lysis buffer (with 50 µg of BSA and 5 µg of sonicated *E. coli* DNA) was subjected to IP using 15 µl of Anti-DYKDDDDK G1 Affinity Resin beads (GenScript) at 4°C for overnight. Five or ten percent of each cell extract was set aside and used to obtain the input DNA samples. IP samples were washed at 4 °C for 5 min with rotation in the following buffers: three times with lysis buffer, once with high salt buffer (50 mM Tris, pH 7.5, 500 mM NaCl, 1 % Triton X-100, 0.1% sodium deoxycholate), once with RIPA buffer (1X TE, 0.25 M LiCl, 0.5 % NP-40, 0.1 % sodium deoxycholate). All lysis and wash buffers contained protease inhibitors. IP samples were eluted with 60 µl of 4 mg/ml 3x FLAG peptide (GlpBio) at 4 °C for 5 hrs with rotation and cross-links were reversed at 65°C for overnight. Samples were treated with RNase A and proteinase K, extracted with phenol-chloroform, precipitated with ethanol, and resuspended in 100 µl of water. The DNA samples were then applied to Hybond-N using a dot blot apparatus, and the membrane was probed with ^32^P-labeled CgTEL-C3 corresponding to three copies of the *C. glabrata* telomeric repeat. Hybridization was performed at 56 °C.

### Structure Modeling of the Cdc13 DBD–junction DNA complex

A model of the *N. glabratus* Cdc13 DNA-binding domain (DBD; UniProt Q6FT40, residues 401–595) bound to a 26-nt junction substrate (5′-AGAAGCATCTGGGTGCTGTGGGGTCT-3′; a 3-bp hairpin stem closed by an AGCA tetraloop, a recessed 5′ end, and a 16-nt telomeric 3′ overhang) was built using the human POT1 DBD–telomeric junction complex (PDB 8SH0) as the template for junction geometry (32). The protein component was the AlphaFold2 model AF-Q6FT40 (mean pLDDT 82.6; low-confidence segments limited to the termini and two surface loops outside the DNA-binding groove) (40). It outperformed MODELLER 10.8 comparative models built on POT1 OB1 alone (GA341 0.15; normalized DOPE +0.89) or together with 1S40 (normalized DOPE +0.09), with a normalized DOPE of −1.41, and it agreed with the experimental *S. cerevisiae* Cdc13 DBD (TM-score 0.79; Cα RMSD 2.43 Å over 171 residues). The DBD was superposed onto POT1 OB1 with US-align (RMSD 3.42 Å over 117 residues). The three junction-proximal base pairs and the recessed 5′ end were taken from the duplex-junction structure 8SH1, and the AGCA loop was taken from 8SH0. Bases were converted to the target sequence on the template backbone. Keeping the POT1 ssDNA path produced extensive steric overlap with the Cdc13 groove, so the overhang was placed on the Cdc13 ssDNA path from 1S40 instead (41), with a two-nucleotide linker spanning the 8.5 Å gap to the duplex.

Complexes were refined in OpenMM 8.6.1 (Amber ff14SB/OL15; OBC2 implicit solvent). Refinement consisted of energy minimization, then 60 ps of Langevin dynamics at 300 K with the protein backbone and hairpin restrained and the ssDNA free, then minimization with restraints released. The refined model contained no protein–DNA interatomic contacts closer than 2.6 Å (compared with >120 severe clashes before refinement). All three stem base pairs kept Watson–Crick geometry (N1–N3, 2.88–3.19 Å), and the protein Cα RMSD from the AlphaFold model was <0.3 Å, so the DNA was accommodated without distorting the fold. The interface buries 1,624 Å² and involves 39 protein residues and 48 polar contacts (≤3.5 Å). Its calculated interaction energy (−214 kcal/mol) was the most favorable of the registers tested. By contrast, ssDNA started on the POT1 path drifted 4.7 Å from it and gave less than half the buried area and interaction energy. The overhang is contacted by aromatic and basic residues typical of OB-fold ssDNA recognition (including Y465, R485, and Y539). Because the force field lacks a terminal 5′-monophosphate, the phosphate was restored at its template position after refinement. In a separate restraint-guided variant (residues 401–415 unrestrained; annealed at 350 K), R411 and R531 reached hydrogen-bonding distance of the 5′-phosphate (3.9 and 3.1 Å) with a core Cα RMSD of 0.32 Å. Because this arrangement was imposed rather than emergent, it is treated as a hypothesis.

### Statistical analysis

Statistical analyses were performed using GraphPad Prism 11.0. Samples were compared and p-values derived using Student’s t-test (two-tailed and unpaired) as implemented in Prism. All C-circle analysis was performed using at least two biological replicates, with each sample analyzed in two separately assays.

## Results

### The loss of *N. glabratus* Cdc13 results in telomere damage mediated by Exo1, Sgs1 and multiple DDR proteins

Because budding yeast Cdc13 is essential, its protection functions have been studied primarily in *S. cerevisiae* using the *cdc13-1 ts* mutant (18). As a complementary approach that avoids the confounding effects of residual Cdc13, we developed a plasmid eviction protocol to analyze *cdc13Δ* null mutants in *N. glabratus*. Strains harboring a chromosomal deletion of *CDC13* were kept viable by a plasmid carrying both *CDC13* and a *URA3* selectable marker (designated *cdc13Δ\**); these strains were grown in non-selective media for ∼8 population doublings to allow loss of the plasmid, and then spotted on both YPD and 5-FOA plates, with growth on 5-FOA (toxic to *URA3+* cells) revealing the phenotype of cells that have lost Cdc13. Deletions in various DDR and DNA repair factors were engineered into *cdc13Δ\** to assess their contributions to growth impairment and telomere damage in the absence of Cdc13. As the point of departure, we targeted the Exo1 and Sgs1 end-resection pathways and the Rad9-dependent DNA damage response pathway, since concurrent inactivation of these pathways is sufficient to confer viability on *S. cerevisiae cdc13-1* cells at non-permissive temperatures (13). The results in *N. glabratus* were largely concordant. None of the strains harboring single or double deletions grew appreciably on 5-FOA medium, whereas the *exo1Δ sgs1Δ rad9Δ* combination mutant grew dramatically better, indicating that these end-resection and DDR pathways contribute independently to the telomere damage and growth defects caused by the loss of Cdc13 (Figure 1A, top two sets of panels).

**Figure 1.**
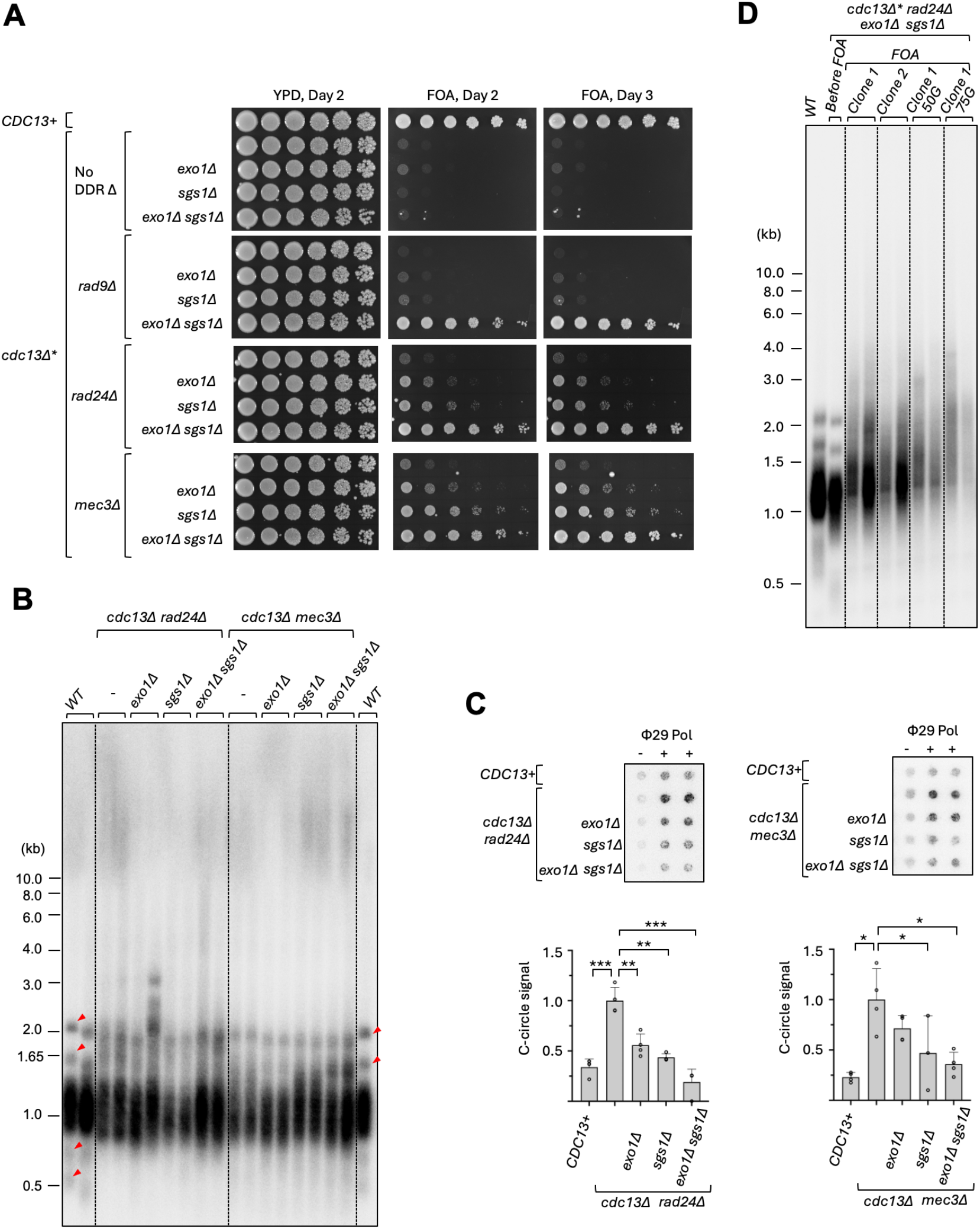
The growth and telomere defects of *N. glabratus cdc13Δ* mutants bearing deletions in DDR and DNA repair genes. **A.** Serial dilutions of the specified, conditional *cdc13Δ* strains were spotted onto YPD and FOA media, and their growth assessed after 2 and 3 days of incubation. **B.** Genomic DNAs from the specified conditional *cdc13Δ* strains were isolated following growth in FOA for ∼5-8 population doublings and subjected to TRF Southern analysis. Genomic DNAs from the parental, wild type strain were analyzed in parallel. Discrete TRF clusters in the wild type samples (which become less distinct in mutant samples owing to increased telomere length heterogeneity) are marked by red arrowheads. **C.** Genomic DNAs from the specified conditional *cdc13Δ* strains were isolated following growth in FOA for ∼5-8 population doublings and subjected to C-circle analysis. Representative C-circle blots are shown at the top and the quantified results plotted at the bottom. **D.** Genomic DNAs were isolated from the wild type and the conditional *cdc13Δ rad24Δ exo1Δ sgs1Δ* strain following growth in the specified media and duration, and then subjected to TRF Southern analysis. Note that over-expression of Cdc13 from the plasmid-borne *CDC13* (with the *PDC1* promoter) in the cdc13Δ* rad24Δ exo1Δ sgs1Δ strain before growth on FOA caused slight telomere shortening.

To gain additional insight into the DDR, we tested components of the 9-1-1 clamp (Mec3) and its clamp loader (Rad24), which have also been implicated in telomere damage. In contrast to *rad9Δ*, *rad24Δ* and *mec3Δ* each synergized with either *exo1Δ* or *sgs1Δ* to confer moderate growth on *cdc13Δ* cells, suggesting that the 9-1-1 clamp and its loader are more dominant DDR factors than Rad9 in this setting (Figure 1A, bottom two sets of panels). Interestingly, however, the *rad24Δ exo1Δ sgs1Δ cdc13Δ* and *mec3Δ exo1Δ sgs1Δ cdc13Δ* mutants grew comparably to *rad9Δ exo1Δ sgs1Δ cdc13Δ*: once both resection pathways are inactivated, deleting any one of the DDR genes improves growth to the same extent. We therefore tested epistasis directly, between *rad24Δ* and *mec3Δ* and between *rad24Δ* and *rad9Δ* (Supp. Figure 1A and 1B). In the context of either *cdc13Δ* or *cdc13Δ exo1Δ sgs1Δ*, neither double mutant improved growth beyond the respective single mutants, supporting the notion that Rad24, Mec3 and Rad9 act in a single DDR pathway in the absence of Cdc13.

Telomere restriction fragment (TRF) Southern analysis of the partially viable *cdc13Δ* strains (those carrying DDR deletions) revealed minimal changes in overall telomere length but substantial increases in heterogeneity, evident as a blurring of the discrete TRF clusters seen in wild type (Figure 1B, WT sample, marked by red arrowheads). Consistent with telomere DNA damage, these mutants also accumulated C-circles, and the amounts of C-circles tracked with the severity of the growth defects (Figure 1C) – both for the series of strains sharing *CDC13* and *RAD24* deletions and for those sharing *CDC13* and *MEC3* deletions.

The initial viability of the *rad24Δ exo1Δ sgs1Δ cdc13Δ* mutant allowed us to passage this strain for ∼75 generations and follow its growth and telomeres. Clones exhibited severe senescence, and TRF Southern analysis revealed progressive loss of telomere content accompanied by a moderate increase in average telomere length during passage (Figure 1D). Because the loss of Cdc13 is expected to abolish telomerase recruitment in this strain, we surmise that the apparent lengthening reflects stochastic, recombination-mediated elongation at a small fraction of chromosome ends, which is ultimately insufficient to counter senescence.

Taken together, these findings identify the long-range resection pathways and a DDR network comprising the DDR clamp, its loader and the Rad9 adaptor as the mediators of telomere DNA damage in Cdc13-deficient cells. Notably, the preferred substrates of both the resection pathways and the DDR clamp loader are ss-dsDNA junctions with recessed 5’ ends, suggesting that the exposure of such structures is a key step in the damage process.

### Disrupting the CST-PP PIC assembly preferentially triggers Exo1-mediated DNA damage

We showed previously that cluster mutations in a surface loop of Ten1 that target the CST-PP-DNA pre-initiation complex (PIC) abrogate CST stimulation of PP activity *in vitro* and, upon over-expression, cause growth and telomere defects *in vivo*. Importantly, these mutations do not affect the telomere localization of Ten1, implying that the protection function intrinsic to CST, independent of the PIC, remains intact. We therefore compared the effects of DDR and DNA repair gene deletions on one of the PIC mutants (*ten1^L34B^*) with their effects on *cdc13Δ*, in order to isolate the protection function of the PIC. Strains harboring combinations of DDR and repair gene deletions were transformed with plasmids over-expressing either wild type Ten1 or *ten1^L34B^* and subjected to growth and telomere analysis. The *ten1^L34B^* mutant protein was expressed at comparable levels in the different strain backgrounds, ruling out spurious effects of differential over-expression (Figure 2A).

**Figure 2.**
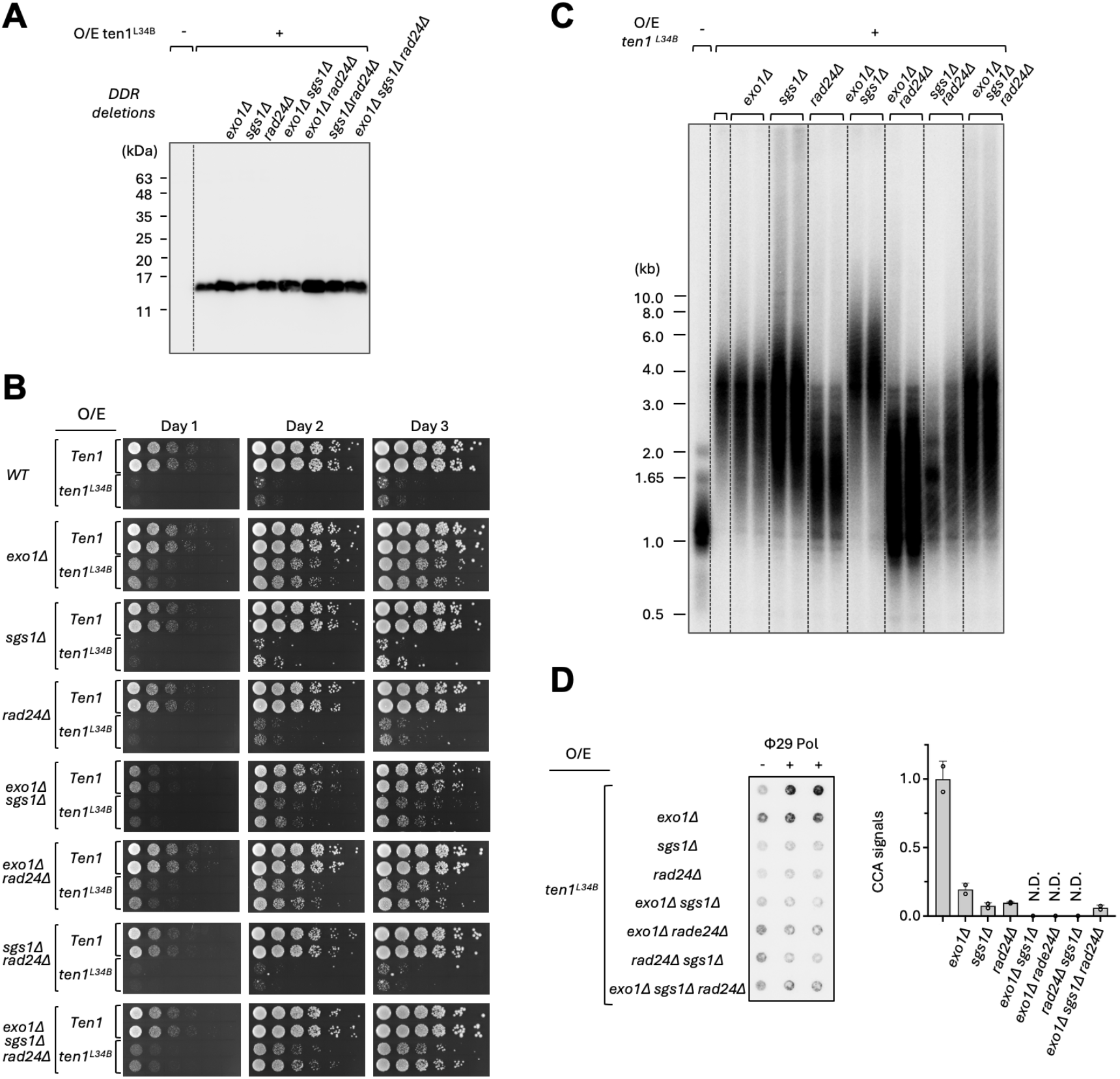
The growth and telomere defects of *N. glabratus* strains that over-express the *ten1^L34B^* mutant protein. **A.** The levels of the *ten1^L34B^* mutant protein in the specified strains that contain the *ten1^L34B^*-overexpression plasmid were analyzed by western blotting. **B.** Serial dilutions of the specified strains harboring plasmids expressing either wild type Ten1 or *ten1^L34B^* were spotted on the SD-ura medium, and their growth assessed after 1, 2 and 3 days of incubation. **C.** The specified strains were transformed with the *ten1^L34B^*-overexpression plasmid. Genomic DNAs were isolated after ∼60 generations of growth and subjected to TRF Southern analysis. **D.** The same genomic DNAs were subjected to C-circle analysis. A representative C-circle blot is shown on the left and the quantified results plotted on the right. N.D. designates samples where quantification yielded negative averages. Note that there is no strict correlation between C-circle levels and growth defects; while C-circles is a marker of telomere damage, the levels of C-circles are regulated by many factors including the DDR and resection factors deleted in these strains.

Growth analysis uncovered clear differences between *cdc13Δ* and *ten1^L34B^*. Whereas *exo1Δ* substantially improved the growth of *ten1^L34B^*, *sgs1Δ* and *rad24Δ* had very modest or no effect (Figure 2B), and combining *exo1Δ* with either *sgs1Δ* or *rad24Δ* or both did not further enhance the growth of strains harboring *ten1^L34B^* relative to Ten1. Thus, in contrast to *cdc13Δ*, the growth deficiency of *ten1^L34B^* is dictated primarily by Exo1. Telomere length analysis, however, revealed an effect of each DDR deletion in *ten1^L34B^*, as well as further differences between the *ten1^L34B^* and *cdc13Δ* series (Figure 2C). As reported previously, and unlike *cdc13Δ*, over-expression of *ten1^L34B^* alone induces dramatic telomere elongation (8). Against this background, *rad24Δ* consistently reduced average telomere lengths, alone or in combination with *exo1Δ* or *sgs1Δ*, indicating that Rad24 contributes to the lengthening. In contrast, *sgs1Δ ten1^L34B^* had longer telomeres than *ten1^L34B^*, and this effect was further enhanced in *sgs1Δ exo1Δ ten1^L34B^*, indicating that Sgs1 and Exo1 instead limit the elongation induced by *ten1^L34B^*. Sgs1 and Exo1 appear to act in a pathway parallel to the Rad24-dependent one that supports elongation: the telomere lengths of *exo1Δ sgs1Δ rad24Δ ten1^L34B^* are intermediate between those of *exo1Δ sgs1Δ ten1^L34B^* and *rad24Δ ten1^L34B^*. C-circle analysis showed that every single and combination DDR deletion reduced the level of this damage-associated structure, consistent with each of these proteins acting on telomere metabolism in the *ten1^L34B^* mutant (Figure 2D).

Together, these findings indicate that the subunits of the CST-PP assembly contribute distinct protection functions. Deletion of *CDC13*, which eliminates both CST and PP, leads to growth defects mediated by multiple resection and DDR factors (Exo1, Sgs1 and Rad24/Mec3/Rad9), whereas disruption of the PIC leads to growth defects mediated primarily by Exo1. The telomere phenotypes of the *ten1^L34B^* mutants nevertheless show that Sgs1 and Rad24 can still modulate telomere metabolism in these cells, albeit without substantially influencing growth.

Notably, we had identified a second *ten1* mutation (*ten1^L34A^*) that also disrupts the CST-PP-DNA PIC but causes telomere elongation without evidence of growth defects or telomere DNA damage (8). We interpret this to mean that partial disruption of the PIC leaves the complex defective in telomerase termination but competent in telomere protection. Consistent with this interpretation, neither *exo1Δ* nor *rad24Δ* substantially affected the telomere elongation induced by *ten1^L34A^* (Supp. Figure 1C), indicating that these repair factors do not act in this lengthening pathway.

### RNase H1 and H2 promote Rad24- and Exo1-dependent telomere DNA damage that is counteracted by Cdc13

As described above, the 9-1-1 clamp loader and the Exo1 nuclease play especially prominent roles in the DDR and end resection triggered in *cdc13Δ* cells. Because both preferentially recognize ss-dsDNA junctions with recessed 5’ ends (42,43), such structures are likely to be exposed and unprotected in Cdc13-deficient cells. One potential source of junctions with recessed 5’ ends is the 5’ end of an Okazaki fragment. Because telomeres are replicated unidirectionally, from proximal origins toward distal ends, these 5’ ends are predicted to lie exclusively on the C-strand. During normal Okazaki fragment maturation, they are likely capped by the RNA primer and its triphosphate moiety, and displacement synthesis by the next, more terminal fragment, combined with flap cleavage and ligation, removes the RNA without ever exposing a ss-dsDNA junction (Supp. Figure 2A). If the process is disrupted, however – by fork stalling or reversal, for instance – the RNA primer may instead be removed by RNase H, exposing the junction. We tested this possibility by asking how deletion of *RNH1* (RNase H1) and/or *RNH201* (the catalytic subunit of RNase H2) affects the growth and telomere defects of *cdc13Δ* and *ten1^L34B^* mutants.

In pilot experiments, neither *rnh1Δ* nor *rnh201Δ* alone affected the growth of *cdc13Δ*, but the *rnh1Δ rnh201Δ* double deletion (abbreviated *rnh1/201ΔΔ*) modestly increased growth, an effect further enhanced in combination with *exo1Δ* (Supp. Figure 2B). We therefore analyzed *rnh1/201ΔΔ* systematically in the context of other DDR and DNA repair gene deletions (Figure 3A). Besides *exo1Δ, rnh1/201ΔΔ* synergized strongly with *rad24Δ* and with *sgs1Δ exo1Δ* to improve the growth of *cdc13Δ*. Strikingly, the improvement conferred by ablating both RNase Hs was minimal in the *exo1Δ rad24Δ cdc13Δ* and *sgs1Δ rad24Δ cdc13Δ* mutants, suggesting that RNase H contributes to telomere damage through pathways that largely overlap with these resection and DDR factors – for example, by generating the structures these proteins recognize. Consistent with this idea, *rnh1/201ΔΔ* also improved the growth of cells over-expressing *ten1^L34B^* (Figure 3B). Expression of *ten1^L34B^* from the moderately strong *HHT2* promoter triggered substantial growth defects that were partially reversed by *rnh1/201ΔΔ*, and expression from the weaker *EGD2* promoter caused milder defects that were likewise alleviated (Figure 3B; compare the sizes of the colonies highlighted by yellow arrowheads). In contrast, *rnh1/201ΔΔ* did not enhance the growth of cells over-expressing *ten1^L34A^*, the PIC mutant that induces telomere elongation rather than DNA damage (Figure 3C), nor did it improve the growth of *ten1^L34B^ exo1Δ* cells (Supp. Figure 2C), implying that the RNase Hs act in the same damage pathway as Exo1 in the *ten1^L34B^* mutant. Together, these findings support the notion that RNase H1 and H2 cooperate with each other, and with the resection and DDR factors, to promote DNA damage in CST-deficient cells.

**Figure 3.**
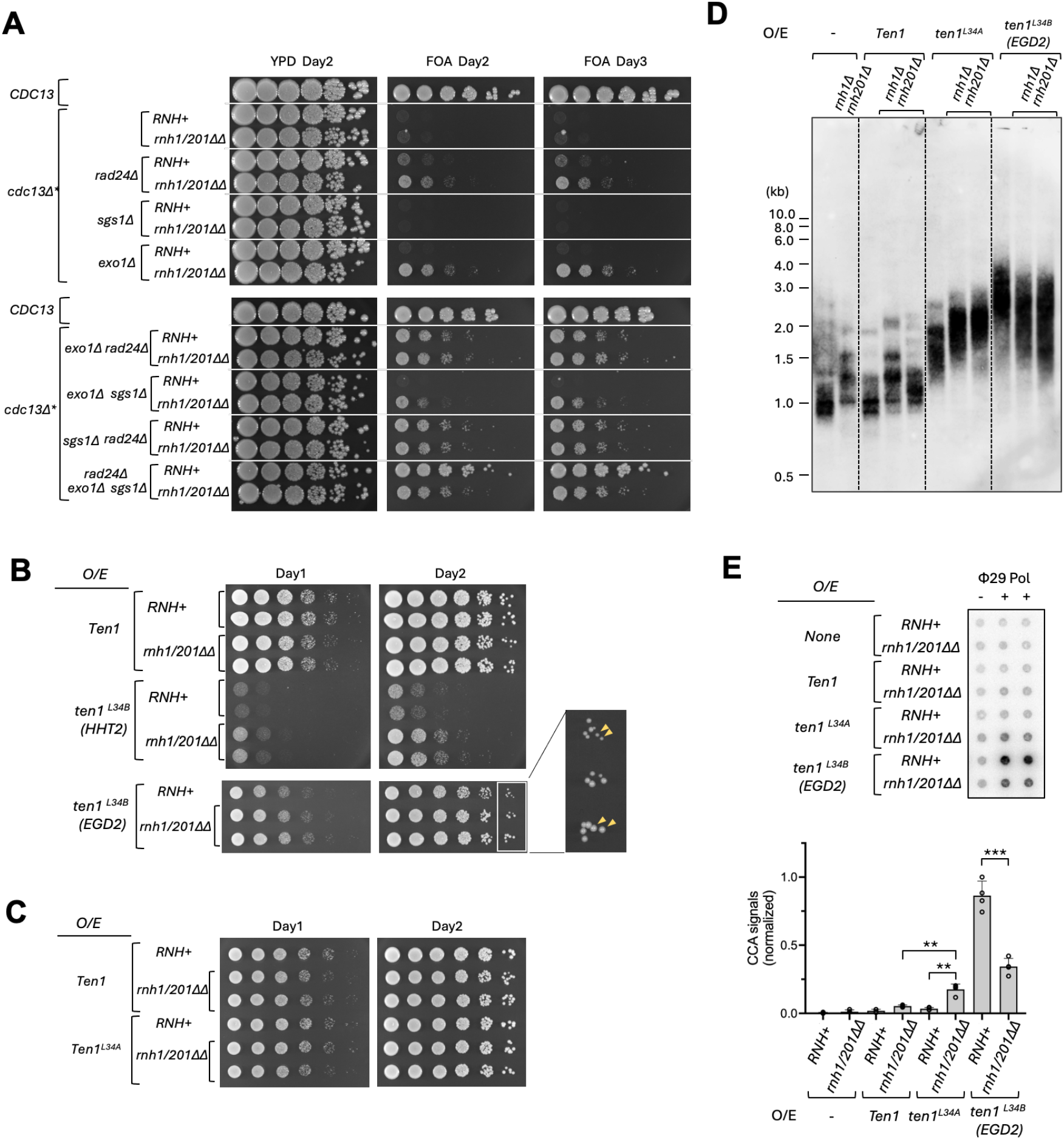
The effects of RNase H1 and H2 deletions on the growth and telomere defects of *CDC13* and *TEN1* mutants. **A.** Serial dilutions of the specified, conditional *cdc13Δ* strains with or without dual deletions of *RNH1* and *RNH201* were spotted onto YPD and FOA media, and their growth assessed after 2 and 3 days of incubation. **B.** The wild type and the *rnh1Δ rnh201Δ* mutant strains were transformed with plasmids over-expressing Ten1 or *ten1^L34B^* (using either the strong *HHT2* or the moderately strong *EGD2* promoter). Serial dilutions of the transformants were spotted onto SD-ura medium, and their growth assessed after 1 and 2 days of incubation. A portion of the day 2 image for the *ten1^L34B^* (EGD2) strains is enlarged to better visualize the colony size differences (marked by yellow arrowheads). **C.** The wild type and the *rnh1Δ rnh201Δ* strains carrying plasmids overexpressing Ten1 or *ten1^L34A^* were analyzed for growth using serial dilutions of cultures spotted onto SD-ura medium and incubated for 1 and 2 days. **D.** Genomic DNAs were isolated from the wild type and the *rnh1Δ rnh201Δ* strains transformed with plasmids over-expressing Ten1, *ten1^L34A^* or *ten1^L34B^* (*EGD2* promoter) after ∼ 75 generations of growth and subjected to TRF Southern analysis. Two independent clones of each *rnh1Δ rnh201Δ* strain were analyzed. The TRFs in this blot were detected using non-radioactive probes followed by chemiluminescence. **E.** The genomic DNAs from the same strains as in D were analyzed for C-circle levels.

We next examined telomeres in strains carrying combinations of *ten1* and RNase H mutations. The *rnh1/201ΔΔ* double deletion alone caused mild telomere elongation and synergized with *ten1^L34A^* to lengthen telomeres further (Figure 3D; Supp. Figure 2D and 2E). These findings are consistent with a recent study in *S. cerevisiae* showing that the RNase Hs contribute to telomere loss by removing the terminal RNA primer after replication (44). In contrast, *rnh1/201ΔΔ* caused mild telomere shortening in cells harboring *ten1^L34B^*, suggesting that these enzymes have different effects at deprotected telomeres. In line with this, *rnh1/201ΔΔ* modestly elevated C-circles in the *ten1^L34A^* mutant but substantially reduced them in *ten1^L34B^* cells (Figure 3E). Overall, the growth and telomere data support the idea that the RNase Hs contribute to telomere DNA damage in CST-deficient cells, possibly by exposing ss-dsDNA junctions, and they also reveal complex effects of these enzymes on the length and structure of protected and deprotected telomeres.

### Identification of Cdc13 residues involved in ss-dsDNA junctional protection

Our data implicate the exposure of ss-dsDNA junctions as an important consequence of Cdc13 loss, suggesting that Cdc13, or CST as a whole, shields such structures from DDR recognition and aberrant repair. This notion echoes a recent study showing that mammalian POT1, the functional counterpart of budding yeast Cdc13, protects the recessed C-strand 5’ end at ss-dsDNA junctions (32). POT1 does so through a surface crater known as the “pothole” – formed by tyrosine and basic residues in an N-terminal loop and the 4^th^ β-strand of its DNA-binding domain – which masks the 5’ terminal nucleotide. Sequence alignment revealed two relatively well-conserved basic residues of Cdc13 (Arg^411^ and Arg^531^) near the equivalent positions, which could perform a similar function (Supp. Figure 3A). Neither residue has been implicated in ssDNA binding by mutagenesis or structural analysis (41,45,46) (Supp. Figure 3B). Moreover, using the DNA from the POT1-ss-dsDNA complex and an AlphaFold model of the *Ng*Cdc13 DBD as templates, we could build a sterically plausible model of a Cdc13-ss-dsDNA complex in which Arg^411^ and Arg^531^ lie on either side of the C-strand 5’ end (Supp. Figure 3C; see Methods for details and for the limitations of the model).

We substituted these basic residues in *N. glabratus* Cdc13 with Glu and, using plasmid shuffle, introduced the single and double mutants (Cdc13^R411E^, Cdc13^R531E^ and Cdc13^RREE^) into the *cdc13Δ\** background. Only the double mutant exhibited substantial growth defects after eviction of the wild type plasmid on FOA (Figure 4A). Southern analysis nevertheless revealed distinct effects of all three alleles on telomere length: Cdc13^R411E^ induced mild elongation, whereas Cdc13^R531E^ and Cdc13^RREE^ produced telomeres that were both longer and more heterogeneous (Figure 4B). After a further ∼75 generations of growth, all three mutants displayed increased length heterogeneity as well as new and variable telomere clusters consistent with subtelomeric recombination (Figure 4C, marked by arrowheads). Together, these results indicate that the putative junction-binding mutants are defective in telomere protection. Western analysis showed that all the mutant proteins accumulate to wild type levels, so the phenotypes cannot be attributed to reduced expression or protein instability (Figure 4D). To probe the mechanism of the growth deficiency, we expressed Cdc13^RREE^ in strains bearing DDR and DNA repair deletions (Figure 4E). Among the single deletions, *rad24Δ* substantially improved growth and *exo1Δ* did so modestly, while *sgs1Δ* had no effect. The double and triple deletion strains mirrored the single ones, with *rad24Δ exo1Δ* and *rad24Δ exo1Δ sgs1Δ* harboring Cdc13^RREE^ showing the best growth relative to the corresponding strains harboring wild type Cdc13. Consistent with increased telomere DNA damage, all the junctional mutants accumulated C-circles, with *cdc13^RREE^* reaching the highest levels (Figure 4F), and in line with the growth suppression, *exo1Δ* and *rad24Δ* each reduced the C-circle accumulation of *cdc13^RREE^*. Deletion of *SGS1*, in contrast, greatly elevated C-circle levels, indicating exacerbated DNA damage. Telomere length analysis in these mutants likewise showed that *exo1Δ* and *rad24Δ* suppress, whereas *sgs1Δ* exacerbates, the telomere aberrations (Figure 4G). We conclude that Cdc13^RREE^ is selectively deficient in protection against Rad24 and, to a lesser extent, Exo1. Notably, despite its slow growth the *cdc13^RREE^* mutant can be passaged extensively (up to 350 generations) and shows modest telomere elongation in later generations (Supp. Figure 4A) – as expected, since the telomerase recruitment function of Cdc13 should be retained in this mutant.

**Figure 4.**
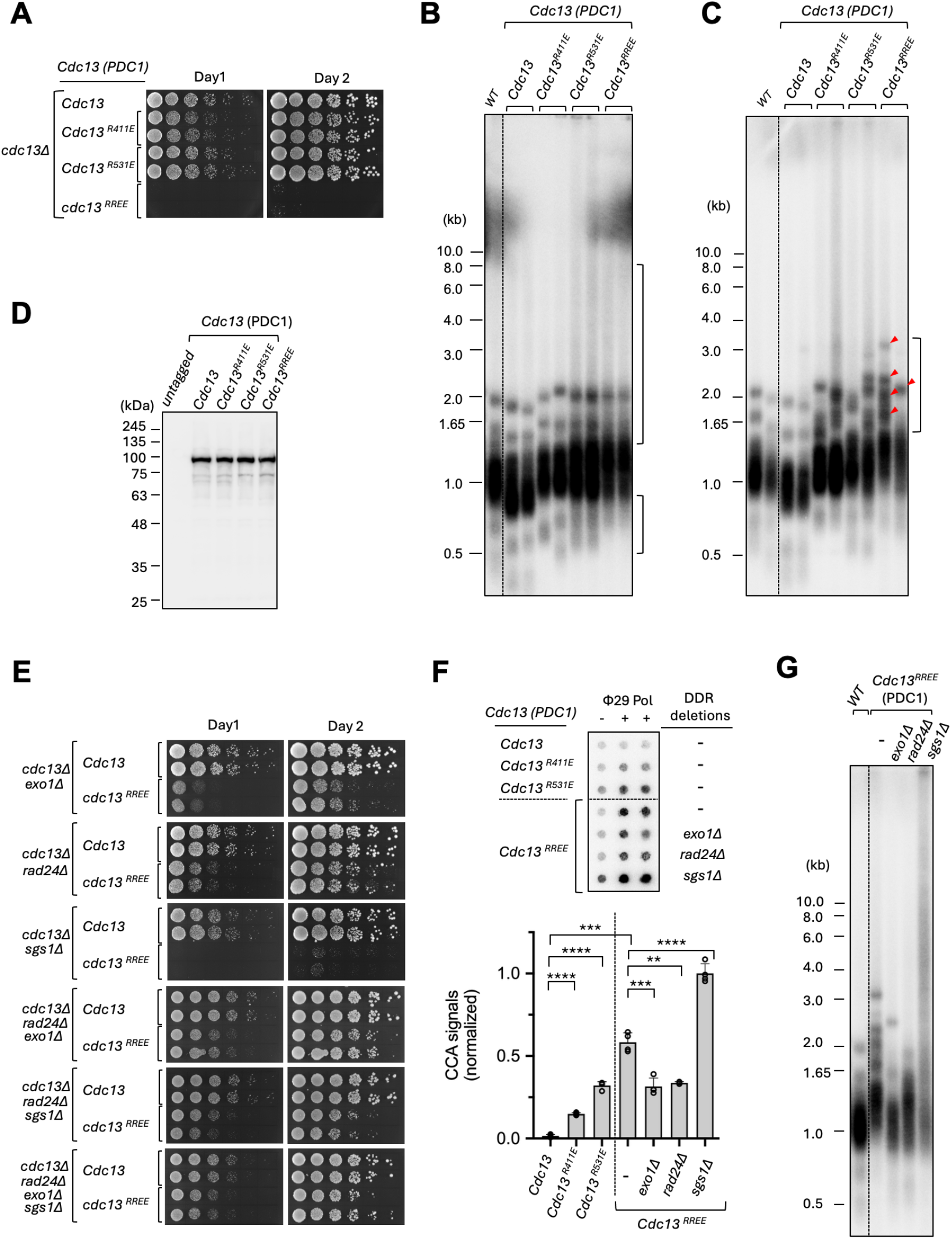
The growth and telomere aberrations of *N. glabratus* strains with a chromosomal *CDC13* deletion and over-expressing Cdc13 mutants with defects in telomere protection. **A.** Serial dilutions of the indicated strains were spotted onto YPD medium and incubated at 30 °C. Images of the plates were collected after 1 and 2 days of incubation. Note that these strains initially contained a wild type *CDC13* on a Ura3-containing plasmid to suppress growth defects triggered by the mutant allele. The strains were cured of the Ura3 plasmid on FOA, and the resulting colonies grown in liquid YPD before being tested in spotting assays. **B.** Genomic DNAs were isolated from the strains in A (two independent clones each) after ∼35 generations of growth without the Ura3 plasmid carrying wild type *CDC13*, and subjected to TRF Southern analysis. **C.** Same as B except that the genomic DNAs were isolated following an additional 75 generations of growth (three streaks for single colonies). **D.** Extracts were prepared from strains harboring the specified Cdc13 mutant, and subjected to Western analysis using antibodies directed against the FLAG_3_ tag fused to each of the mutants. **E.** Serial dilutions of the indicated strains were spotted onto YPD medium and incubated at 30 °C. Images of the plates were collected after 1 and 2 days of incubation. The cells in liquid cultures used for spotting have undergone ∼35 generations of growth without the Ura3 plasmid carrying wild type *CDC13*. **F.** Genomic DNAs from the indicated strains were subjected to C-circle analysis. A representative blot is shown at the top and the quantified results plotted at the bottom. **G.** Genomic DNAs from the indicated strains were subjected to TRF Southern analysis.

In the plasmid shuffle system the Cdc13 mutants were expressed from the strong *PDC1* promoter. We also determined the consequences of expressing Cdc13^R411E^, Cdc13^R531E^ and Cdc13^RREE^ from the native promoter, by integrating each mutant (with a C-terminal FLAG_3_ tag) at the native locus in *cdc13Δ\**. In this configuration the *cdc13^R531E^* strain exhibited mild growth defects and *cdc13^RREE^* was inviable, so the greater viability of the same alleles expressed from *PDC1* indicates that the defects of the junctional mutants are partially alleviated by over-expression (Supp. Figure 4B). As in the over-expression strains, the growth defects of the integrated *cdc13^R531E^* and *cdc13^RREE^* alleles were greatly suppressed by *rad24Δ exo1Δ sgs1Δ* (Supp. Figure 4B, bottom panel), confirming loss of protection against these proteins. In line with the growth defects, the re-integrated *cdc13^R531E^* strain had elongated, heterogeneous telomeres and high levels of C-circles, all substantially ameliorated in the *rad24Δ exo1Δ sgs1Δ* background (Supp. Figure 4C and 4D). We conclude that the putative junctional mutants are primarily defective in protection against Rad24- and Exo1-mediated telomere damage, and that this damage is partially suppressed by over-expression: *cdc13^RREE^* is inviable when integrated at the native locus, yet the same allele expressed from *PDC1* supports viability for more than 300 generations.

### The Cdc13 junctional mutants display altered telomere C-strand 5’ ends

The POT1 pothole mutants harbor altered telomere C-strand 5’ ends, in line with abnormal C-strand degradation (32). Given the apparent similarities between Cdc13 and POT1 in junctional protection, we investigated the effects of the Cdc13 mutations on *N. glabratus* C-strand 5’ ends. Mapping preferred C-strand 5’ ends has been challenging in the standard budding and fission yeasts because of telomere repeat sequence heterogeneity; the only budding or fission yeast characterized in this regard is *Naumovozyma castellii*, which harbors an 8-bp regular repeat and a preferred C-strand nucleotide accounting for ∼30% of 5’ ends (47). In pilot experiments we analyzed the C-strand 5’ ends of the BG14 parental strain using a well-established ligation-PCR protocol together with *N. glabratus*-specific primers (Figure 5A; Supp. Figure 5A and 5B) (38,39). Genomic DNA was first ligated to a mixture of telorette oligonucleotides, each carrying an anchor segment (Teltail) and an annealing segment, so that they can be juxtaposed to all potential C-strand 5’ ends (Supp. Figure 5A). Using the ligated DNA as template, two subtelomeric forward primers (F1 and F4), each annealing to a subset of chromosome ends, generated the predicted heterogeneous PCR products in conjunction with the Teltail reverse primer (Supp. Figure 5B). We then carried out PCR on genomic DNAs ligated to individual telorette oligonucleotides. Reactions with either forward primer revealed preferential positioning of the C-strand 5’ ends at positions 6 and 13 (Supp. Figure 5C and 5D), with the weakest signals consistently at positions 1 and 10, roughly midway between the two peaks. The *N. glabratus* telomere C-strand therefore terminates preferentially at positions 6 and 13 of the 16-bp regular repeat.

**Figure 5.**
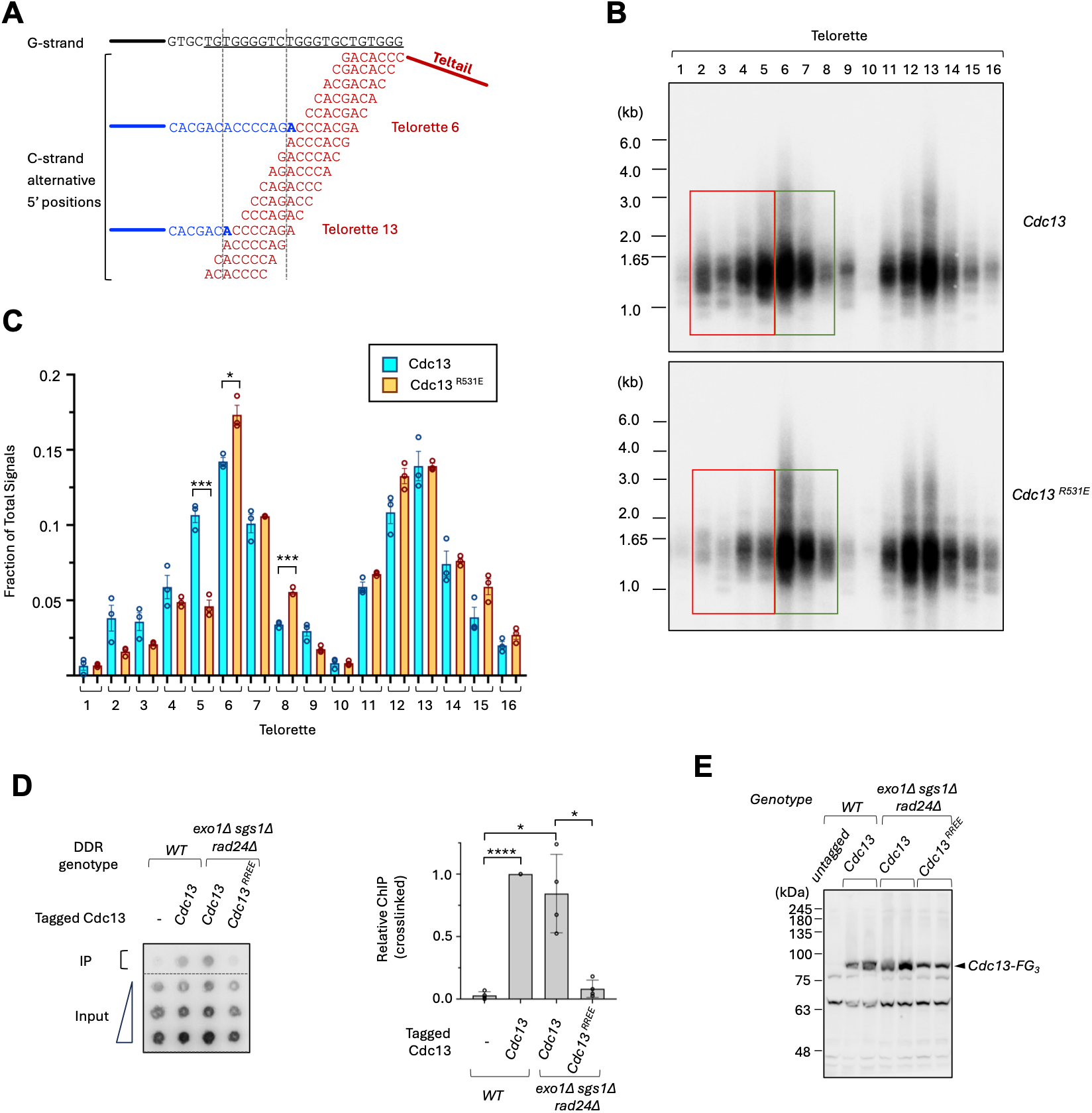
Alteration of the telomere C-strand 5’ end distribution by a *cdc13* protection mutant; Reduced association of a *cdc13* protection mutant with telomere DNA *in vivo*. **A.** Schematic illustration of the ligation-PCR method used to map the *N. glabratus* telomere C-strand 5’ ends. The 3’ portions of the telorette oligonucleotides, engineered for ligation to all possible C-strand 5’ ends, are shown in red. A telomere G-strand sequence as well as the two most common C-strand terminal sequences (ligatable to telorette 6 and telorette 13) are displayed. **B.** Representative blots of the ligation-PCR analysis for telomere C-strand 5’ ends are shown for the strains with wild type Cdc13 and *cdc13^R531E^* integrated at the native chromosome locus. The differences in signals surrounding the peak at position 6 between the Cdc13 and *cdc13^R531E^* telomeres are highlighted by red and green boxes. **C.** The fractions of telomere C-strands that ligate to specific telorettes are quantified and plotted for the Cdc13 and *cdc13^R531E^* strains. Data (mean +/- S.D.) are from three independent experiments. **D.** The indicated strains were analyzed for Cdc13-telomere association *in vivo* using chromatin immunoprecipitation. A representative blot showing input and precipitated signals (crosslinked samples) is shown on the left and the quantified results plotted on the right. Data (mean +/- S.D.) are from 4 independent experiments. **E.** Western analysis of Cdc13 levels in the strains used for the chromatin immunoprecipitation assays.

We then assessed the effect of a junction-protection mutation on this distribution, using the re-integrated wild type *CDC13* and *cdc13^R531E^* strains. To resolve differences more accurately, we reduced the PCR cycle number and quantified the products by Southern hybridization. As in the pilot experiments, the wild type strain showed two major peaks, centered on positions 6 and 13 (Figure 5B and 5C). Both peaks were retained in the *cdc13^R531E^* mutant, but the intensity distribution shifted, most noticeably across the peak and shoulders surrounding position 6: signals from positions 2 – 5 were consistently reduced and those from positions 6 – 8 increased (Figure 5B, compare the red and green boxes in the top and bottom panels), with several individual positions reaching statistical significance (Figure 5C). The peak surrounding position 13 showed a mild overall increase, but no individual position reached significance there. These results suggest that defects in junctional protection by Cdc13 alter the telomere C-strand 5’ ends, while leaving the two preferred positions intact – implying that other factors also contribute to positioning these ends.

Next, we asked whether the junctional protection defect compromises the telomere localization of Cdc13 *in vivo*. Because the re-integrated *cdc13^RREE^* mutant is inviable, we analyzed the *rad24Δ exo1Δ sgs1Δ cdc13^RREE^* strain instead. Chromatin immunoprecipitation (ChIP) revealed substantial enrichment of wild type Cdc13 on telomere DNA in both the wild type and the *rad24Δ exo1Δ sgs1Δ* background (Figure 5D), whereas very little association was detected between *cdc13^RREE^* and telomere DNA in the *rad24Δ exo1Δ sgs1Δ* background, even though western analysis revealed comparable levels of the mutant protein (Figure 5E). Junctional binding may therefore contribute to the stable association of Cdc13 with telomere DNA. One caveat is that *rad24Δ exo1Δ sgs1Δ cdc13^RREE^* grows poorly and its lysates consistently contain lower levels of input DNA (Figure 5E), which may itself affect the efficiency of ChIP.

### Rif1 mediates telomere protection against Rad24 and Exo1 in the context of CST deficiency

The data presented so far underscore the importance of Cdc13 and CST-PP in protecting ss-dsDNA junctions and antagonizing aberrant DDR recognition and repair. In this respect Cdc13 resembles Rif1, another telomere regulatory factor with junctional binding and protection activity (28). In *S. cerevisiae*, Rif1 supports the function of CST in telomere capping and, like CST, regulates telomere length by modulating telomerase activity (27,29,48). The precise physical and functional relationships between Rif1 and CST, and the linkage between their protection and length regulatory functions, nevertheless remain unclear. We examined these issues in *N. glabratus* by combining *rif1Δ* with CST mutations.

First, we introduced *rif1Δ* into a set of conditional *cdc13Δ* strains carrying resection and DDR deletions. Loss of Rif1 further reduced the residual growth of *cdc13Δ*, *cdc13Δ rad24Δ* and *cdc13Δ rad24Δ sgs1Δ* but had little or no effect on *cdc13Δ rad24Δ exo1Δ* and *cdc13Δ rad24Δ exo1Δ sgs1Δ* (Figure 6A). These results are consistent with Rif1 supporting the capping function of CST, and further suggest that Rif1 antagonizes primarily Exo1- and Rad24-mediated damage. The protection defect of *rif1Δ* thus resembles that of *cdc13^RREE^*, which also manifests primarily Rad24- and Exo1-dependent damage (Figures 4E – 4G). To examine this further, we analyzed *rif1Δ* in strains harboring *cdc13^RREE^* together with DDR deletions and found the same pattern as in the corresponding *cdc13Δ* strains (Supp. Figure 6A): *rif1Δ* worsened the growth of *cdc13^RREE^*, *cdc13^RREE^ rad24Δ* and *cdc13^RREE^ rad24Δ sgs1Δ* but had little effect on *cdc13^RREE^ rad24Δ exo1Δ* and *cdc13^RREE^ rad24Δ exo1Δ sgs1Δ.* We conclude that Rif1 is important for supporting the protection functions compromised by the *cdc13^RREE^* mutations.

**Figure 6.**
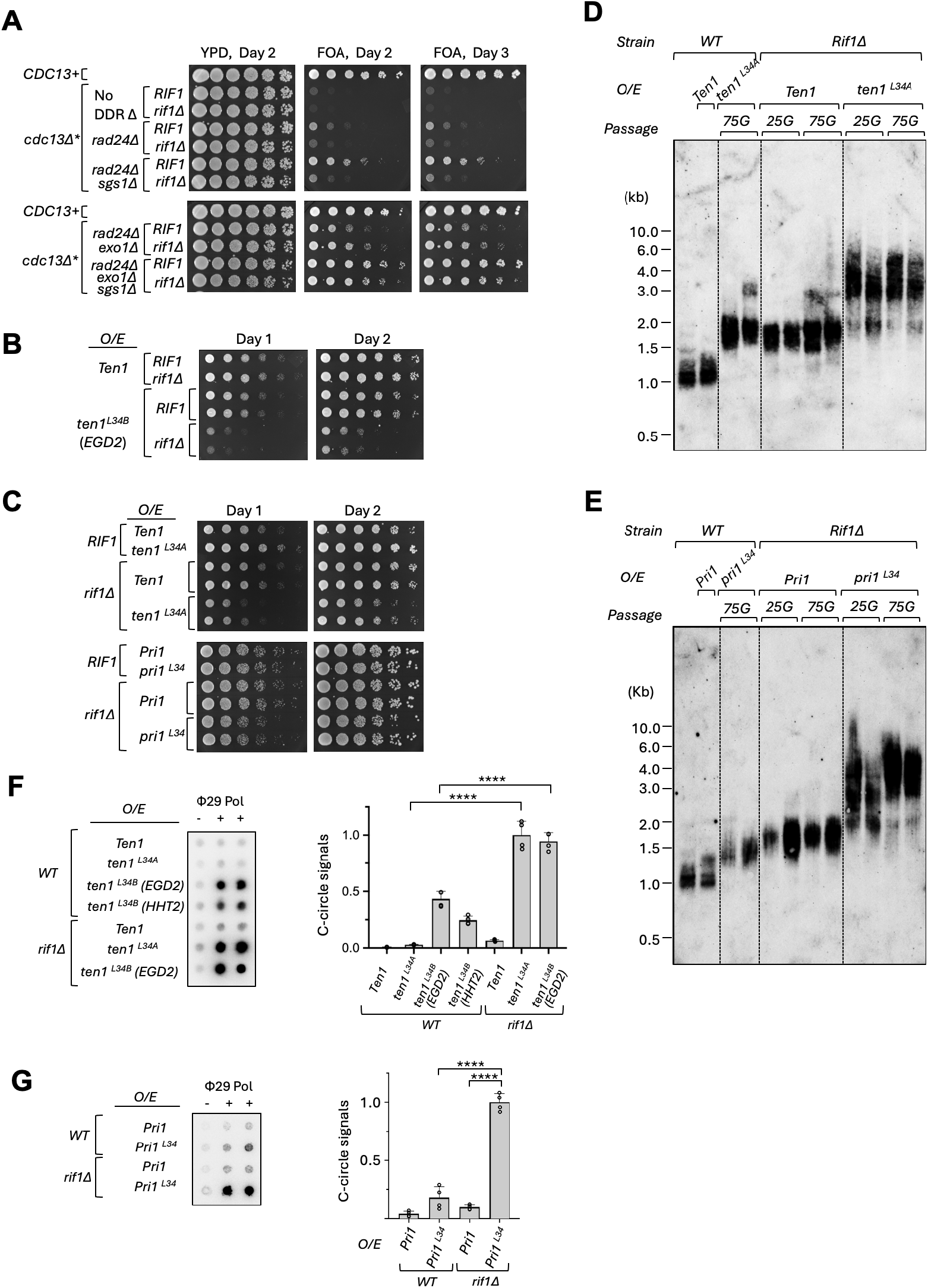
The effects of *RIF1* deletion on the growth and telomere defects of *CDC13* and *TEN1* mutants. **A.** Serial dilutions of the specified, conditional *cdc13Δ* strains with or without *RIF1* deletion were spotted onto YPD and FOA media, and their growth assessed after 2 and 3 days of incubation. **B.** The wild type and *rif1Δ* strains were transformed with plasmids over-expressing Ten1 or *ten1^L34B^* (*EGD2* promoter). Serial dilutions of the transformants were spotted onto SD-ura medium, and their growth assessed after 1 and 2 days of incubation. **C.** The wild type and *rif1Δ* strains carrying plasmids overexpressing Ten1, Ten1^L34A^, Pri1, or Pri1^L34^ were analyzed for growth using serial dilutions of cultures spotted onto SD-ura medium and incubated for 1 and 2 days. **D.** Genomic DNAs were isolated from the wild type and the *rif1Δ* strains transformed with plasmids over-expressing Ten1 or Ten1^L34A^ after the specified generations of growth and subjected to TRF Southern analysis. Two independent clones of each strain were analyzed. The TRFs in this blot were detected using non-radioactive probes followed by chemiluminescence. **E.** Genomic DNAs were isolated from the wild type and the *rif1Δ* strains transformed with plasmids over-expressing Pri1 or Pri1^L34^ after the specified generations of growth and subjected to TRF Southern analysis. Two independent clones of each strain were analyzed. The TRFs in this blot were detected using non-radioactive probes followed by chemiluminescence. **F.** Genomic DNAs from the indicated strains were subjected to C-circle analysis. A representative blot is shown on the left and the quantified results plotted on the right. **G.** Same as F, for the strains shown in E.

Next, we tested *RIF1* deletion in the context of two *ten1* mutants that disrupt the CST-PP-DNA PIC, and observed notable effects on both. While moderate over-expression of *ten1^L34B^* (EGD2-*ten1^L34B^*) alone caused only a very mild growth defect, *rif1Δ* EGD2-*ten1^L34B^* showed profound growth deficiency (Figure 6B). Further elevating *ten1^L34B^* expression (HHT2-*ten1^L34B^)* caused apparent synthetic lethality with *rif1Δ*: no colonies could be recovered from the selection plate when *rif1Δ* was transformed with the HHT2-*ten1^L34B^* plasmid (Supp. Figure 6B). Likewise, while *ten1^L34A^* alone showed no growth defect, as was also true of *rif1Δ*, the *rif1Δ ten1^L34A^* double mutant clearly grew more slowly than either single mutant (Figure 6C, top panels). We obtained the same result with *pri1^L34^*, another mutant that disrupts the PIC and causes telomere elongation like *ten1^L34A^*: neither *pri1^L34^* nor *rif1Δ* is growth defective, yet *rif1Δ pri1^L34^* showed notable growth retardation relative to each single mutant (Figure 6C, bottom panel). Hence Rif1 supports the protection function not only of Cdc13 and CST but of the entire CST-PP-DNA assembly.

We then analyzed telomere lengths and C-circle levels in these combination mutants. We found that *ten1^L34A^* and *pri1^L34^* each induced progressive telomere elongation, as shown previously (8), while *rif1Δ* alone produced stably elongated telomeres (Figure 6D, *cf. rif1Δ Ten1* at 25G and 75G; Figure 6E, *cf. rif1Δ Pri1* at 25G and 75G). Both *rif1Δ ten1^L34A^* and *rif1Δ pri1^L34^* showed more drastic elongation than their respective single mutants, and each double mutant lengthened progressively, as the corresponding *ten1^L34A^* and *pri1^L34^* single mutants do (Figure 6D and 6E). These results support the notion that *rif1Δ* lengthens telomeres by a mechanism distinct from that of *ten1^L34A^* and *pri1^L34^*. In addition, *rif1Δ* synergized separately with *ten1^L34A^*, *ten1^L34B^* and *pri1^L34^* to increase C-circle levels (Figure 6F and 6G), again supporting the idea that Rif1 cooperates with the CST-PP PIC assembly in telomere protection.

To gain insight into the mechanism of this cooperation, we assessed potential physical association between Cdc13-FG_3_ and Rif1-Myc_5_ by co-immunoprecipitation from strains expressing both tagged proteins. No Rif1-Myc_5_ was detectable in pull-down samples enriched for Cdc13-FG_3_, indicating that the two proteins do not form a stable complex in cell extracts (Supp. Figure 6C). We also examined a potential role for Rev7, which is physically linked to Rif1 in mammalian cells and implicated in regulating non-homologous end joining in budding yeast (49,50). However, *rev7Δ* had no effect on the growth or telomere phenotypes of any *CDC13* or *TEN1* mutant tested, arguing against a role for Rev7 in telomere protection (Supp. Figure 6D – 6F).

## Discussion

In this study we identified multiple DDR and end resection factors that mediate telomere damage in cells defective in the CST-PP assembly. Our results highlight the exposure of ss-dsDNA junctions (SDJ) as a critical step in aberrant telomere repair and, conversely, establish junctional protection as a critical activity of CST-PP (Figure 7). They also offer insight into how SDJ become exposed and how CST-PP and related factors protect this structure. We discuss the implications below.

**Figure 7.**
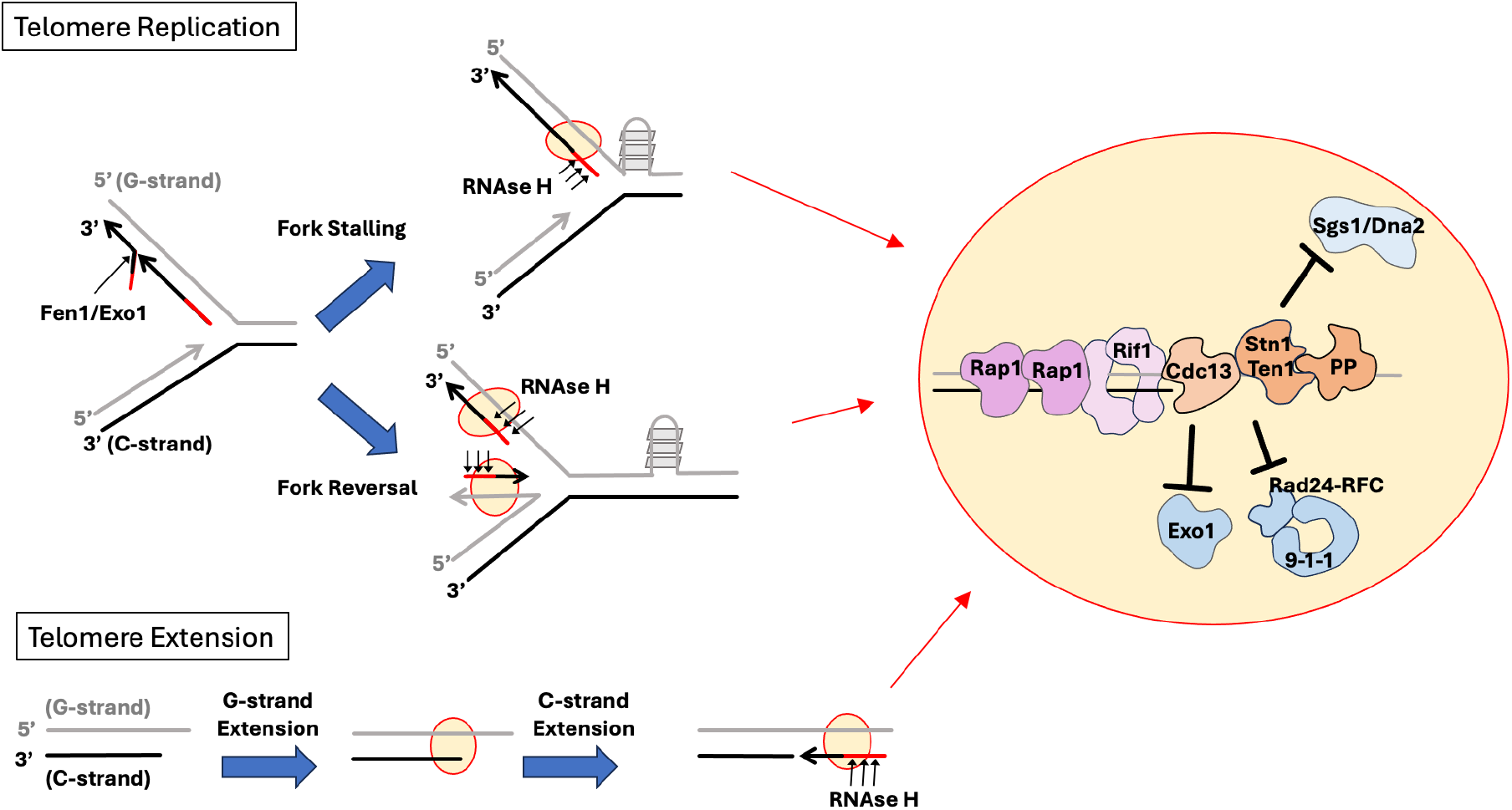
Model for replication-associated telomere DNA damage and protection. (Left) The mechanisms that can create ss-dsDNA junctions during telomere replication and during telomere extension by telomerase and CST-PP are illustrated. The potential sites of junctional DNA that may trigger DDR and aberrant repair are highlighted by red circles shaded in yellow. (Right) The pathways responsible for telomere DNA damage and the proposed telomere protection assembly are illustrated.

### The activation of multiple repair pathways at CST-deficient telomeres

Using a conditional *cdc13Δ* strain in combination with deletions in DDR and DNA repair genes, we showed that the loss of Cdc13 causes growth and telomere abnormalities dependent on a network of DDR factors (Rad9, Rad24 and Mec3) and on end resection proteins (Exo1 and Sgs1). Among these, the DDR clamp and its loader (Mec3 and Rad24) appear especially critical for growth arrest: deleting either gene alone modestly improves proliferation in the absence of Cdc13, and *mec3Δ* and *rad24Δ* each displays strong synergy with *exo1Δ* or *sgs1Δ* in improving the growth of *cdc13Δ* cells. Interestingly, although *RAD9* functions in the same pathway as *RAD24* and *MEC3* by epistasis analysis, *rad9Δ* alone has no effect and synergizes poorly with either *exo1Δ* or *sgs1Δ*. This difference may reflect Rad9 acting as a DDR adaptor that amplifies damage signaling downstream of the clamp and clamp loader. Like *rad9Δ*, eliminating Exo1 or Sgs1 (or both) failed to improve *cdc13Δ* growth, suggesting that these factors also act relatively late in the damage process; the synergy of *exo1Δ* with *rad24Δ/mec3Δ*, and of *sgs1Δ* with *rad24Δ/mec3Δ*, nevertheless indicates that the two resection pathways contribute independently to the growth deficiency. Overall, the response of *N. glabratus cdc13Δ* cells to DNA repair gene deletions closely resembles earlier analyses of the *S. cerevisiae cdc13-1 ts* mutant, supporting conservation of the pathways responsible for growth arrest (13,51). Consistent with telomere damage driving the growth deficiency, C-circles were elevated in *cdc13Δ* cells and their levels tracked inversely with proliferation (Figure 1C). Telomere Southern analysis, however, revealed little change in average telomere length and only a modest increase in heterogeneity, suggesting that *cdc13Δ* cells do not engage in the high levels of telomere recombination characteristic of post-senescent survivors.

In contrast to *cdc13Δ* cells, which lack the entire CST-PP assembly, the *ten1^L34B^* mutant, disrupted only for PIC formation, experiences growth defects mediated primarily by Exo1. Deleting *EXO1* alone substantially improved the growth of cells over-expressing *ten1^L34B^*, whereas deleting *SGS1* or *RAD24* had no effect (Figure 2B), and the growth of *ten1^L34B^ exo1Δ* was not improved further by deleting *SGS1* or *RAD24* or both. The full CST-PP PIC assembly thus appears to be especially important for protecting cells against Exo1. Telomere Southern analysis, on the other hand, revealed distinct effects of *exo1Δ*, *sgs1Δ* and *rad24Δ* on the telomere lengths of the *ten1^L34B^* mutant, indicating that each factor can gain access to the deprotected telomere and modulate the lengthening and/or shortening pathways. Together these results underscore the importance of the PIC in fully protecting telomeres and show that Cdc13 has protection functions beyond the PIC. The selective protection afforded by the PIC against Exo1 also raises a mechanistic question. The anti-resection effect of CST-PP has generally been attributed to fill-in synthesis; if that were the sole mechanism, however, it is unclear why CST-PP should not protect comparably against Sgs1. One possibility is that the PIC also provides a physical block to Exo1 that is independent of fill-in synthesis.

### RNase Hs promote telomere DNA damage in the context of CST deficiency and play multi-faceted roles at protected and de-protected telomeres

Among the DDR and resection factors responsible for the growth and telomere defects of *cdc13Δ* cells, both the DDR clamp and Exo1 preferentially recognize ss-dsDNA junctions with recessed 5’ ends, suggesting that this structure is a major trigger of DNA damage and is normally protected by CST-PP. Because the protection function of Cdc13 appears to be especially critical during DNA replication (12,21), we considered mechanisms that could generate such junctions in S-phase. Drawing on what is known about Okazaki fragment maturation, telomere replication fork stalling and RNA degradation activities, we hypothesized that premature removal of Okazaki fragment RNA primers by RNase H1 and H2 at stalled forks exposes ss-dsDNA junctions to damage recognition and aberrant repair. In support of this idea, RNase H1 and H2 jointly reduce the viability of *cdc13* and *ten1* mutants and exacerbate telomere aberrations in these mutants (Figure 3 and Supp. Figure 2). The combined effects of *RNH1* and *RNH201* deletion on Cdc13-deficient cells are greatly reduced when the cells also harbor combinations of *exo1Δ*, *rad24Δ* and *sgs1Δ*, indicating that the RNase Hs act in pathways overlapping those of these DDR and resection factors. Two further observations support a role for RNase H in primer removal at telomeres. First, RNase H1 and H2 were recently shown to remove the terminal RNA primer from lagging strand telomeres after replication in *S. cerevisiae* (44). Second, *S. pombe* RNase H2 promotes replication fork degradation by removing the RNA primer at reversed forks (52).

Beyond promoting telomere DNA damage, RNase H1 and H2 also regulate telomere length differently at protected and uncapped telomeres. When telomeres are protected, deletion of *RNH1* and *RNH201* increased average telomere length (Figure 3, strains over-expressing Ten1 and Ten1^L34A^), consistent with the recent observation that these nucleases promote telomere shortening by removing the terminal RNA primer (44). When telomeres are uncapped, in contrast, the same deletions decreased the lengths of abnormally elongated telomeres (Figure 3, strains over-expressing Ten1^L34B^), consistent with the nucleases promoting the exposure of damaged DNA. Together these findings highlight the impact of RNase H on telomere metabolism through RNA primer removal. Previous studies have attributed the effects of RNase H at telomeres primarily to activity on TERRA in R-loops (53,54). While our data do not exclude this possibility, it is unclear why removing TERRA from an R-loop should expose junctional DNA unless the junction exists in the first place, with its ssDNA in hybrid with TERRA. Further work will be needed to delineate the respective roles of TERRA and RNA primers in the damage pathways suppressed by CST.

### Protection of ss-dsDNA junctions by Cdc13: molecular mechanisms, similarities to POT1 and evolutionary implications

Starting from the hypothesis that Cdc13 protects ss-dsDNA junctions by a mechanism similar to that of POT1, we identified two conserved basic residues in *Ng*Cdc13 (Arg^411^ and Arg^531^) that, when mutated to acidic residues, caused profound growth and telomere defects. Several lines of evidence implicate these residues in the binding and protection of ss-dsDNA junctions. First, in multiple sequence alignments each lies close to the POT1 residues implicated in binding the C-strand 5’ phosphate at an SDJ (Supp. Figure 3). Second, the *S. cerevisiae* Cdc13 Lys^504^ residue, equivalent to *Ng*Cdc13 Arg^411^, was recently implicated in junction binding (55). Third, a sterically plausible model of a Cdc13-ss-dsDNA complex, built from the SDJ of the POT1 complex and the AlphaFold *Ng*Cdc13 model, accommodates both residues on either side of the C-strand 5’ end, although their contact with the 5’ phosphate had to be imposed by restraints rather than emerging spontaneously (see Methods). Fourth, mutating Arg^411^ and Arg^531^ to Glu caused growth defects, telomere aberrations and altered C-strand 5’ end distribution, all consistent with loss of junctional protection. Finally, the telomere damage in these mutants is caused primarily by Rad24 and Exo1, both of which target ss-dsDNA junctions (42,43,56,57). Conclusive validation of the hypothesis will nonetheless require high resolution structural information.

The junctional protection activity we uncovered for Cdc13 has evolutionary implications and raises questions about similar activities in related proteins. Although the evolutionary relationship between Cdc13 and POT1 has long been controversial, recent analysis suggests that Cdc13 evolved from POT1 through a series of domain re-organizations (58), and the apparent conservation of junction binding and protection in both proteins further supports a close relationship between the two families. POT1 and Cdc13 also share the function of regulating telomere G- and C-strand synthesis, and they do so in analogous ways. For example, POT1 helps recruit and activate telomerase through its partner TPP1, whereas Cdc13 recruits telomerase by binding Est1 and activates it through a network of interactions involving Est3, the TPP1 ortholog (59). In addition, POT1 recruits the CTC1-STN1-TEN1 complex to promote Pol α-mediated C-strand synthesis, whereas Cdc13 recruits the Stn1-Ten1 subcomplex to the same effect (15,60). Together, these structural and functional considerations strongly support a common ancestry. Given the prevalence of POT1 in metazoans and fungi and the restricted presence of Cdc13 in Saccharomycotina yeasts, Cdc13 most likely arose much more recently from a POT1 family member (58). A corollary is that Cdc13 should not be regarded as the ortholog of mammalian or fungal CTC1, despite sharing the C designation in the CST complex (7): unlike Cdc13, CTC1 is a larger protein with seven OB folds that functions primarily in C-strand synthesis and has no role in telomerase recruitment (7). Whether CTC1 possesses junction binding or protection activity is an interesting open question. Mammalian CTC1 antagonizes BLM- and EXO1-mediated resection at DSBs (17), but whether this entails a pothole-like mechanism remains unclear.

### The cooperation between Rif1 and CST-PP in junctional protection at telomeres

Rif1 was originally discovered as a yeast protein that regulates telomere length (48). Subsequent studies showed that it also has a significant role in telomere protection when CST function is compromised (27), and that it possesses junction-binding activity, encasing the duplex portion of the substrate and thus offering a plausible mechanism for protection (28). Our analysis of Rif1 in *N. glabratus* is consistent with these ideas and provides additional insight into how Rif1 cooperates with CST-PP to regulate telomere length and protection. First, the protection function of Rif1 appears to be normally redundant with CST-PP but to become prominent when CST-PP is compromised: whereas *rif1Δ* alone exhibits no telomere protection defect, a variety of CST-PP mutations – in Cdc13, Ten1 and Pri1 – synergize with loss of Rif1 to cause telomere damage and growth defects. Second, in support of junctional protection as the basis of that function, Rif1 preferentially antagonizes Rad24 and Exo1.

Third, Rif1 evidently regulates telomere length by a mechanism distinct from that of CST-PP, since combining *rif1Δ* with *ten1^L34A^* or *pri1^L34^* dramatically accelerated elongation; this is also consistent with current models in which Rif1 affects the loading of telomerase whereas CST-PP removes telomerase after G-strand extension (8–10,61). The protection activity of Rif1, by contrast, is likely to be exercised at the same step as that of CST-PP, given their functional redundancy. An attractive possibility is that when ss-dsDNA junctions are exposed during replication or C-strand fill-in synthesis, CST-PP occupies the ssDNA portion and protects the C-strand 5’ end while Rif1 encircles the dsDNA portion. CST-PP alone suffices for protection, but when it is compromised – by defective PIC assembly, for example – Rif1 could partially block DDR and resection at the junction (Figure 7). Notably, and as in a previous analysis in *S. cerevisiae*, we could not detect a stable association between Rif1 and Cdc13, suggesting that their functional cooperation does not depend on a direct physical interaction (27). A further implication of these observations is that although Rif1 and CST-PP each regulate both telomere length and telomere protection, they do so in different relationships to one another. In length regulation they act at separate steps: Rif1 acts upstream of telomerase, by reducing its access to telomeres, whereas CST-PP acts downstream, by limiting the extent of G-strand extension. In protection, by contrast, CST-PP and Rif1 likely act together, forming a joint assembly on junctional DNA that arises either as a replication intermediate or during telomere extension by telomerase and CST-PP (Figure 7). In summary, ss-dsDNA junctions with recessed 5’ ends are a major trigger of telomere DNA damage and require protection by the CST-PP assembly as well as by Rif1. This damage is enhanced by RNase H, most likely through removal of the RNA primers of Okazaki fragments. Collectively, our findings on CST-PP, RNase H and Rif1 illustrate both the dangers posed by the structural intermediates of telomere replication and elongation and the multifaceted mechanisms that cells have evolved to manage them.

## Data Availability

The data generated and/or analyzed during this study are included in the paper and the Supplementary Data, or are available from the corresponding author upon reasonable request.

## Acknowledgments

We thank members of the Lue lab for discussions. The authors declare no competing interests.

