## Supplemental Figures 1-6 and Tables 1-2 for "CST–Polα/primase, backed up by Rif1, protects telomeric ss-dsDNA junctions from damage recognition and resection"

**Supplementary Figures 1 – 6**

**Supplementary Tables 1 – 2**

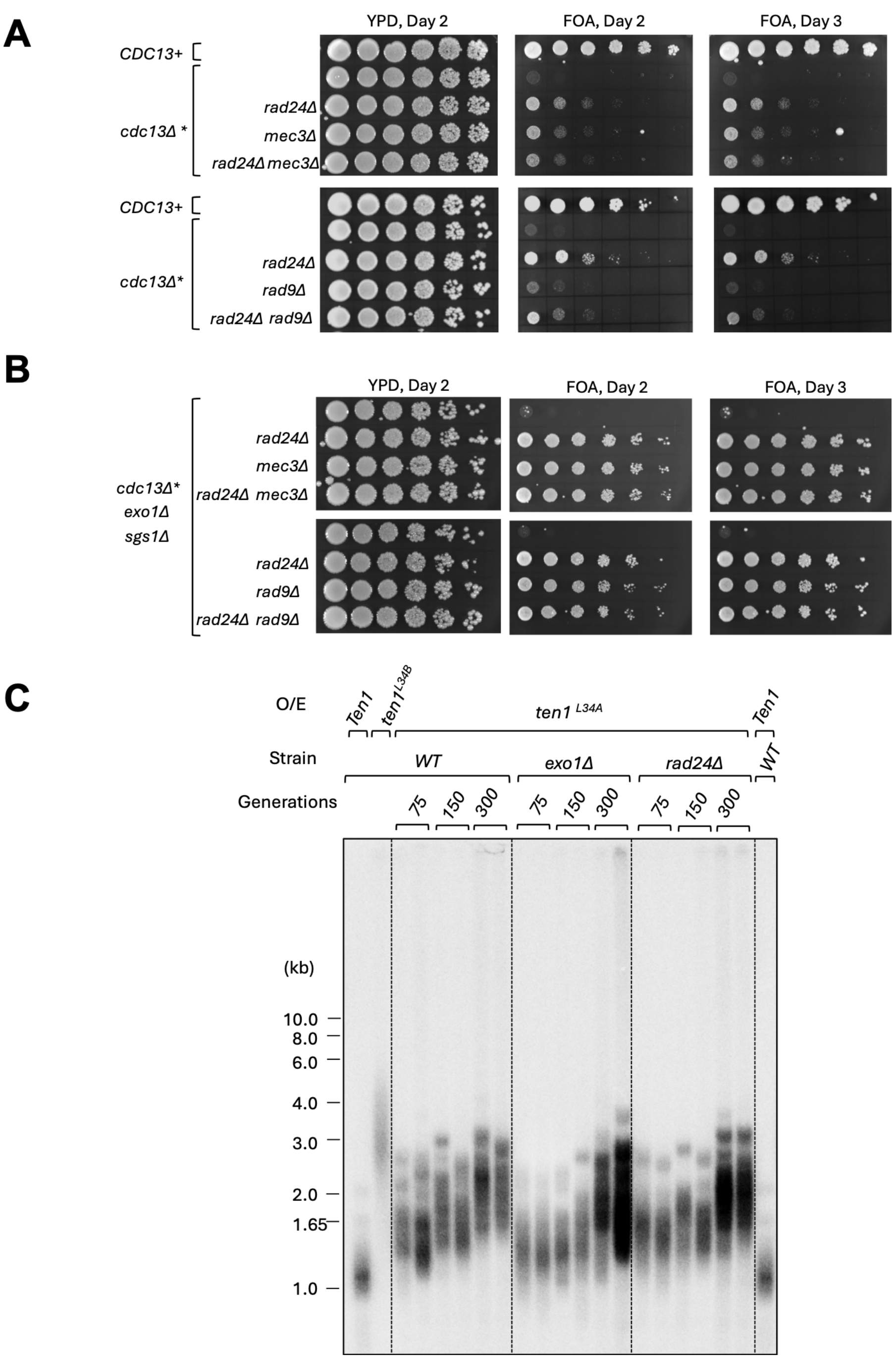

**Supp. Fig. 1. Epistasis between DDR factors in promoting growth defects in *cdc13*Δ cells; failure of *EXO1* and *RAD24* deletions to suppress the telomere elongation induced by ten1$^{L34A}$**

**A. and B.** Serial dilutions of the specified, conditional *cdc13*Δ strains were spotted onto YPD and FOA media, and their growth assessed after 2 and 3 days of incubation. A: Epistasis of DDR factors in the *cdc13*Δ* background**.** B: Epistasis of DDR factors in the *cdc13*Δ* *exo1*Δ *sgs1*Δ background.

**C.** Genomic DNAs from the specified strains over-expressing wild type or mutant Ten1 were isolated after the indicated numbers of generations of growth and subjected to TRF Southern analysis.

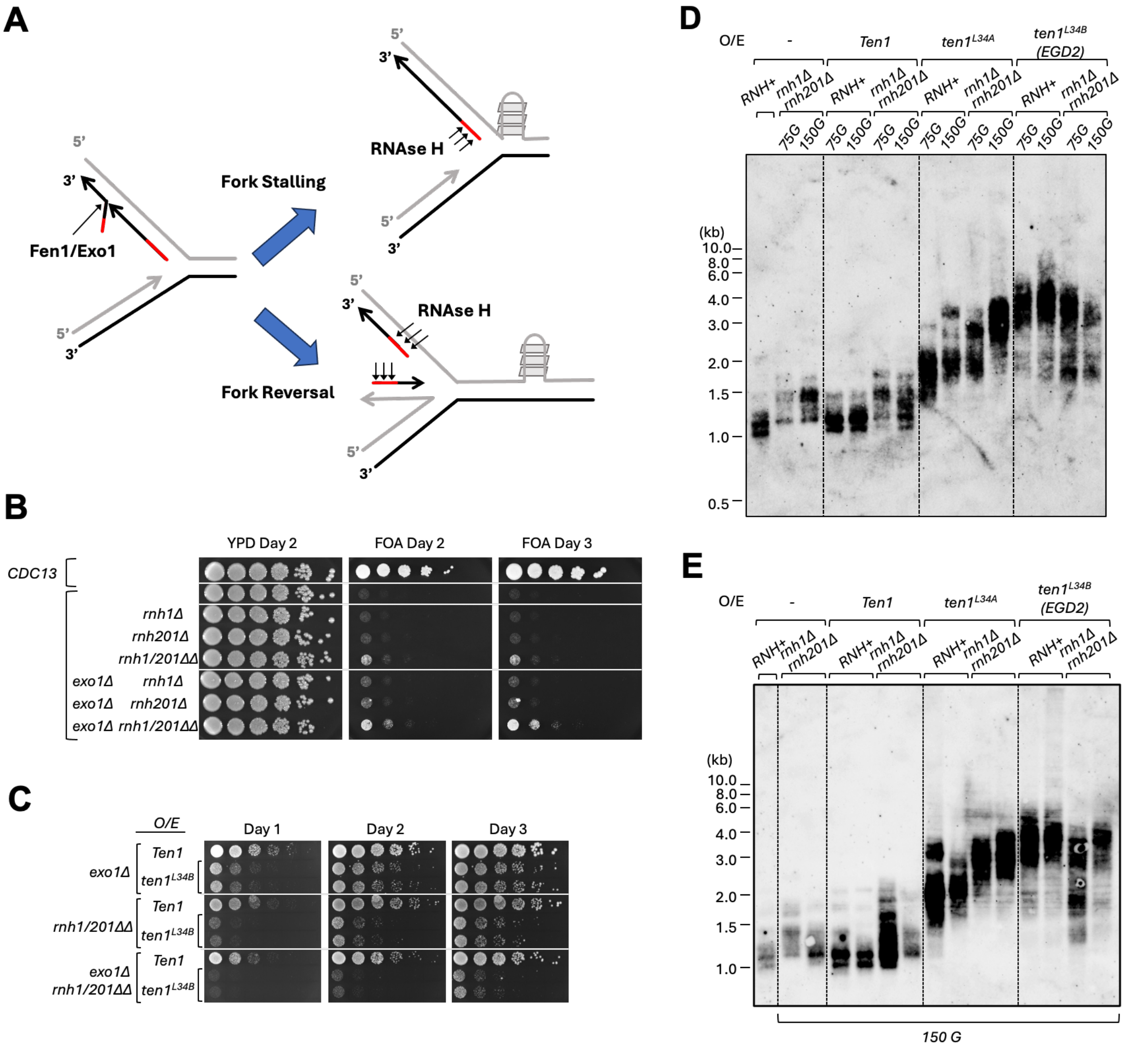

**Supp. Fig. 2. The effects of RNase H1 and H2 deletions on the growth and telomere defects of *CDC13* and *TEN1* mutants**

**A.** Model for the aberrant degradation of RNA primers by RNase H at stalled or reversed forks.

**B.** Serial dilutions of the specified, conditional *cdc13*Δ strains with or without dual deletions of *RNH1* and *RNH201* were spotted onto YPD and FOA media, and their growth assessed after 2 and 3 days of incubation.

**C.** Strains with combinations of *exo1*Δ and *rnh1/201*ΔΔ were transformed with plasmids over-expressing Ten1 or *ten1*$^{L34B}$ (using the strong *HHT2* promoter). Serial dilutions of the transformants were spotted onto SD-ura medium, and their growth assessed after 1, 2 and 3 days of incubation.

**D.** Genomic DNAs were isolated from the wild type and the *rnh1*Δ *rnh201*Δ strains transformed with plasmids over-expressing Ten1, *ten1*$^{L34A}$ or *ten1*$^{L34B}$ (*EGD2* promoter) after ~ 75 or 150 generations of growth and subjected to TRF Southern analysis. The TRFs in this blot were detected using non-radioactive probes followed by chemiluminescence.

**E.** Genomic DNAs were isolated from the wild type and the *rnh1*Δ *rnh201*Δ strains transformed with plasmids over-expressing Ten1, *ten1*$^{L34A}$ or *ten1*$^{L34B}$ (*EGD2* promoter) after ~ 150 generations of growth and subjected to TRF Southern analysis. Two independent clones of each strain were analyzed to assess clonal variability. The TRFs in this blot were detected using non-radioactive probes followed by chemiluminescence.

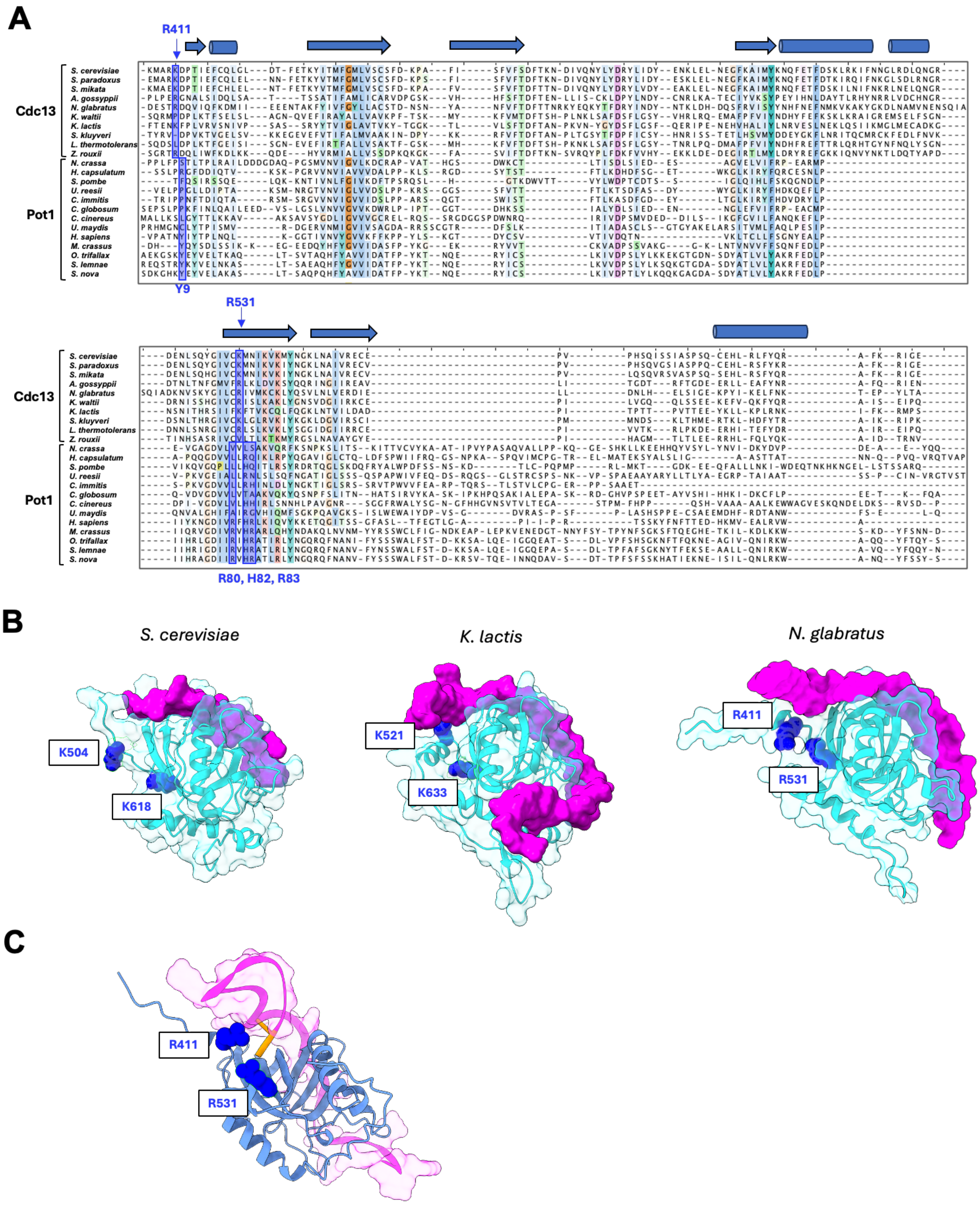

**Supp. Fig. 3. Multiple sequence alignment of Cdc13 and POT1 DNA-binding domains; experimental and computational models of Cdc13-DNA complexes**

**A.** A multiple sequence alignment of the Cdc13 and POT1 DNA-binding domains is shown, with the conserved secondary structural elements displayed on top. The positions corresponding to the proposed junction-binding residues in Cdc13 and the experimentally determined junction-binding residues in POT1 are highlighted by blue-shaded columns.
**B.** Crystal structures and a computational model of Cdc13-ssDNA complexes are displayed, highlighting the lack of contact between the putative junction-binding residues and the ssDNA. The DNA-binding domains, ssDNA and junction-binding residues are shown in cyan (ribbon), magenta (surface) and blue (spheres), respectively. The *S. cerevisiae* and *K. lactis* structures are experimentally determined (PDB: 1S40 and PDB: 6LBR); the *N. glabratus* structure is based on AlphaFold model AF-Q6FT40.
**C.** A computational model of *N. glabratus* Cdc13 DBD bound to a ss-dsDNA junction. The DBD and DNA are shown in light blue and magenta, respectively. The 5' nucleotide of the C-strand is highlighted in orange, and the proposed junction-binding residues are shown as blue spheres. The variant shown is the restraint-guided model, in which $Arg^{411}$ and $Arg^{531}$ were driven to within hydrogen bonding distance of the 5' phosphate; this arrangement was imposed rather than emergent (see Methods).

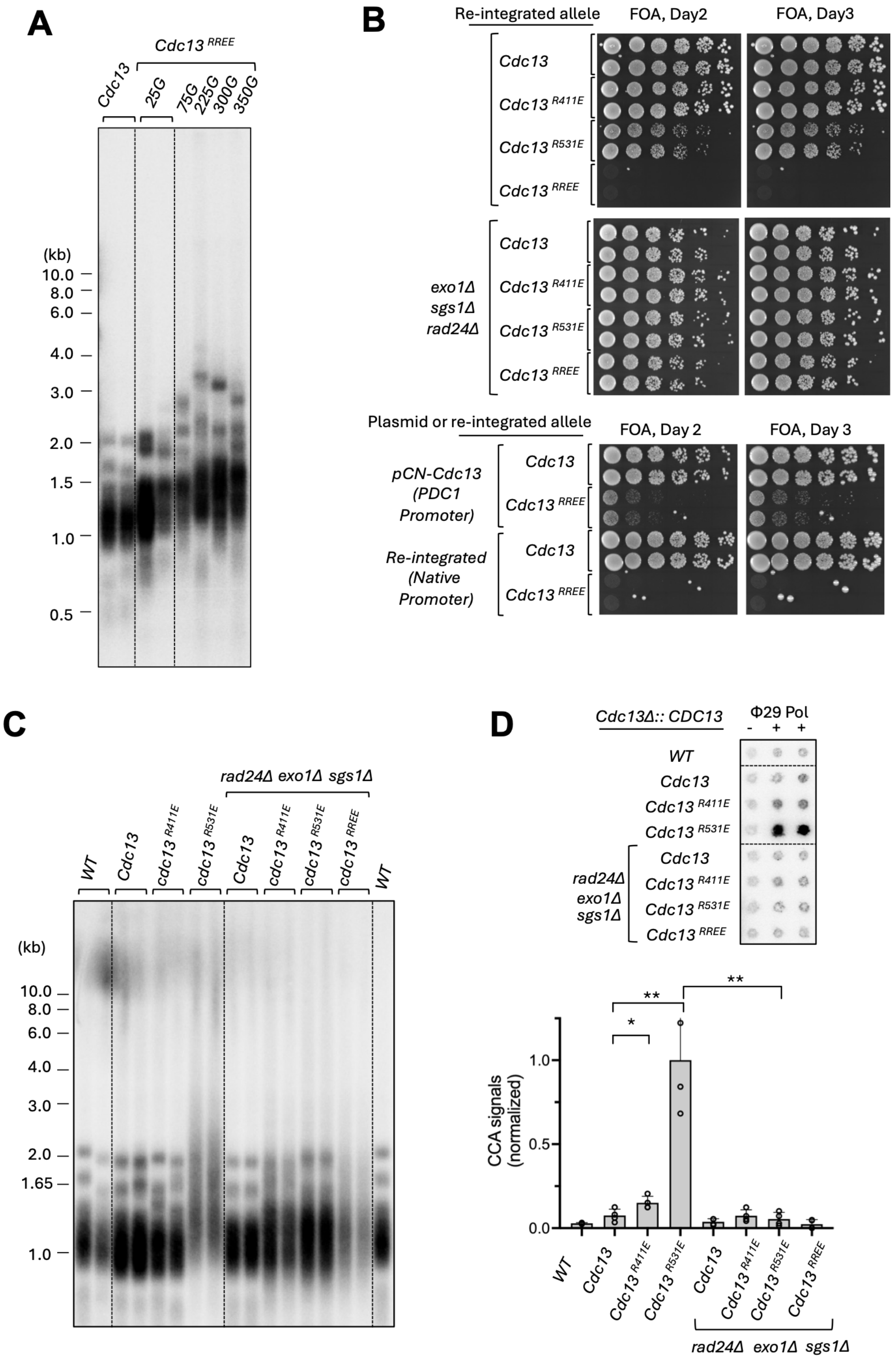

**Supp. Fig. 4. The growth and telomere aberrations of *N. glabratus* strains with chromosomally integrated *CDC13* mutants harboring defects in telomere protection**

**A.** Genomic DNAs were isolated from the strain harboring Cdc13$^{RREE}$ expressed from the strong *PDC1* promoter after the indicated numbers of generations of growth and subjected to TRF Southern analysis.

**B.** Serial dilutions of the indicated strains were spotted onto FOA medium and incubated at 30 °C. Images of the plates were collected after 2 and 3 days of incubation. Note that these strains initially contained a

wild type *CDC13* on a Ura3-containing plasmid to suppress the growth defects triggered by the mutant allele. The strains were first grown in YPD for ~ 8 population doublings to allow for loss of the Ura3 plasmid and then tested in spotting assays.
**C.** Genomic DNAs were isolated from the specified strains (all with chromosomally integrated *CDC13* mutants) after ~35 generations of growth without the Ura3 plasmid carrying wild type *CDC13*, and subjected to TRF Southern analysis.
**D.** The same genomic DNAs were subjected to C-circle analysis. A representative blot is shown at the top and the quantified results plotted at the bottom.

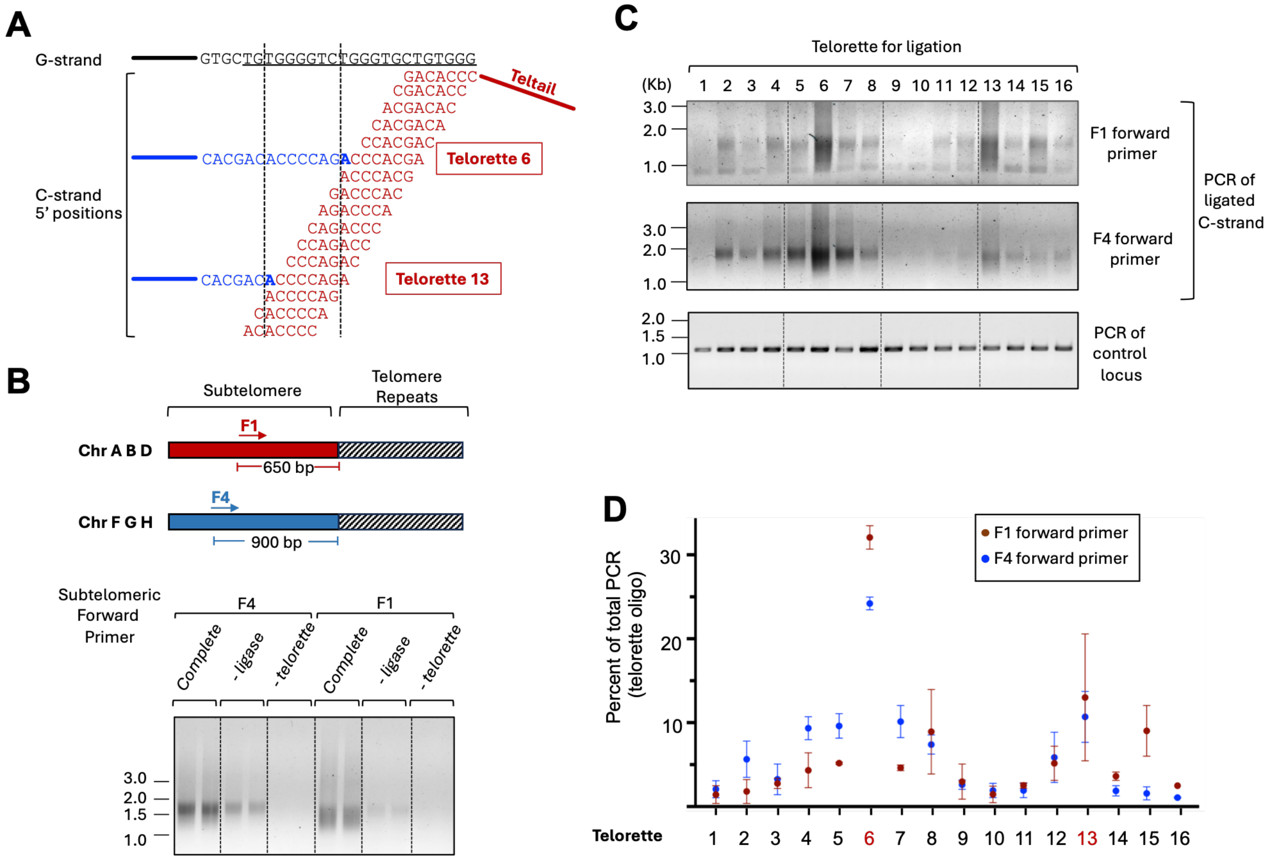

**Supp. Fig. 5. Mapping of the telomere C-strand 5' end distribution in *N. glabratus* wild type cells**
**A.** Schematic illustration of the ligation-PCR method used to map the *N. glabratus* telomere C-strand 5' ends. The 3' portions of the telorette oligonucleotides, engineered for ligation to all possible C-strand 5' ends, are shown in red. A telomere G-strand sequence as well as the two most common C-strand terminal sequences (ligatable to telorette 6 and telorette 13) are displayed.
**B.** (Top) Illustration of the annealing positions of the F1 and F4 forward primers, engineered to detect different groups of *N. glabratus* subtelomeres. (Bottom) Pilot ligation-PCR assays using mixtures of telorette oligonucleotides and omitting specific components (e.g., DNA ligase) are shown.
**C.** Representative gels of the ligation-PCR analysis for telomere C-strand 5' ends are shown for assays using the F1 and F4 forward primers. Amplification of a comparably sized *CDC13* ORF region was used as a control for the quality of the telorette-ligated DNA templates.
**D.** The fractions of telomere C-strands that ligate to specific telorettes are quantified and plotted for assays using the F1 and F4 forward primers.  Data (mean +/- S.D.) are from three independent experiments.

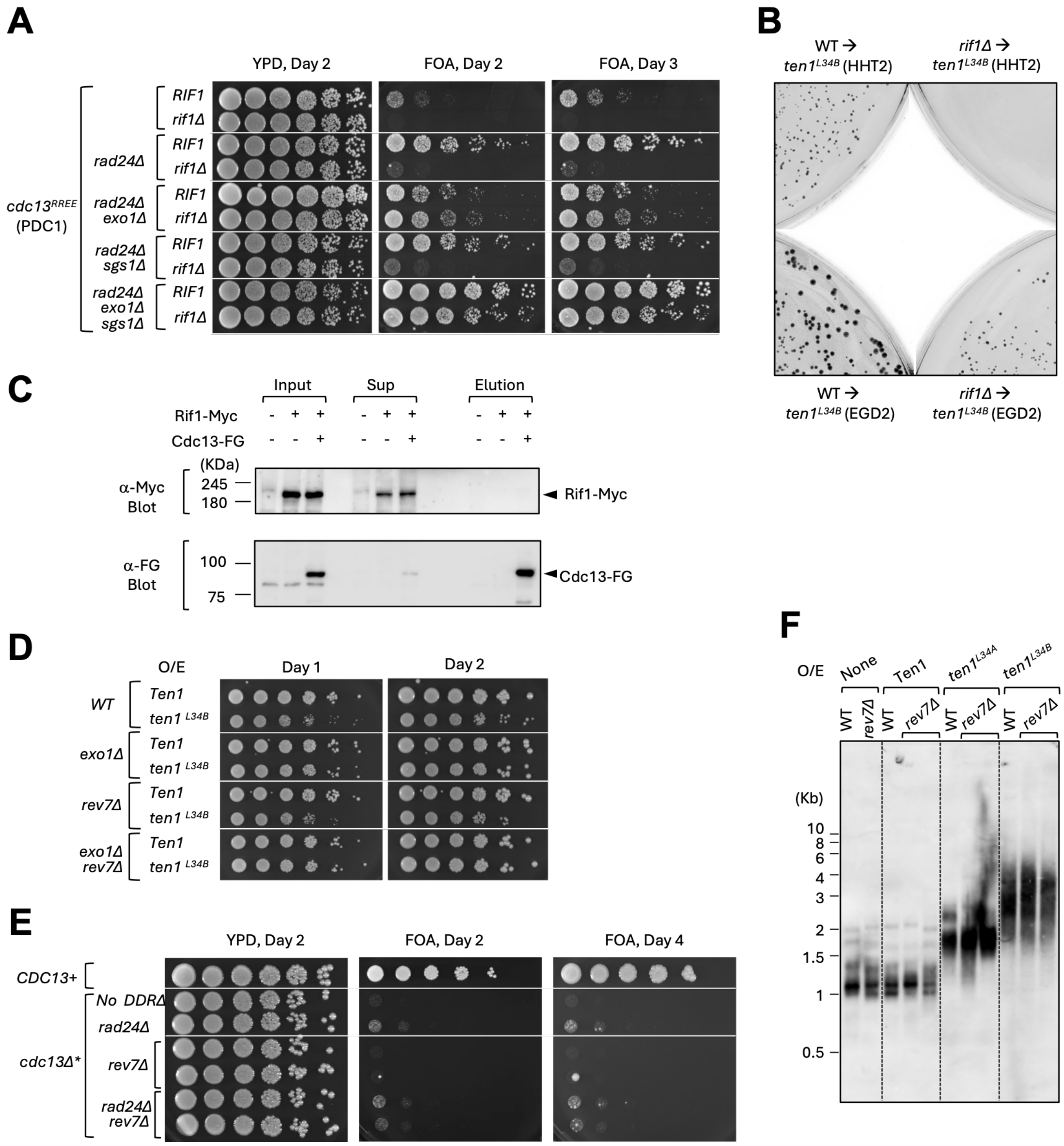

**Supp. Fig. 6. The effects of *RIF1* and *REV7* deletions on the growth and telomere defects of *CDC13* and *TEN1* mutants; lack of a strong physical association between Cdc13 and Rif1 in cell extracts**

**A.** Serial dilutions of the specified, conditional *cdc13*Δ strains with or without *RIF1* deletion were spotted onto YPD and FOA media, and their growth assessed after 2 and 3 days of incubation. Note that all strains express the $Cdc13^{RREE}$ mutant from the strong *PDC1* promoter.

**B.** The wild type and *rif1*Δ strains were each transformed with plasmids expressing $Ten1^{L34B}$ from either the strong *HHT2* promoter or the moderately strong *EGD2* promoter. The numbers and sizes of the transformants were compared after 3 days of incubation.

**C.** Extracts were prepared from strains harboring the indicated $FG_3$-tagged Cdc13 and $Myc_5$-tagged Rif1, and subjected to anti-FG antibody pull down and FLAG peptide elution. The input, supernatant and elution fractions were analyzed by anti-FG and anti-Myc western.

**D.** The indicated strains were transformed with plasmids over-expressing Ten1 or *ten1*$^{L34B}$ (*EGD2* promoter). Serial dilutions of the transformants were spotted onto SD-ura medium, and their growth assessed after 1 and 2 days of incubation.

**E.** The indicated conditional *cdc13*Δ strains with additional *rev7*Δ and/or *rad24*Δ were tested for growth on YPD and FOA media. Serial dilutions of the clones were assessed for growth after 2 and 4 days of incubation.
**F.** Genomic DNAs were isolated from the wild type and the *rev7*Δ strains transformed with plasmids over-expressing Ten1, $Ten1^{L34A}$ and $Ten1^{L34B}$ and subjected to TRF Southern analysis. Two independent clones of each *rev7*Δ strain were analyzed. The TRFs in this blot were detected using non-radioactive probes followed by chemiluminescence.

**Supplementary Table 1. *N. glabratus* strains used in this study**

| **Alias** | ***Relevant Genotype*** | **Parental Strain** | **Reference** |
|---|---|---|---|
| *BG14* | *ura3Δ::Tn903* $G418^R$ | BG2 | Cormack and Falkow, 1999 |
| *exo1Δ* | *exo1Δ::loxP* [a] | BG14 | Calugaru et al., 2025 |
| *rad24Δ* | *rad24Δ::loxP* | BG14 | Calugaru et al., 2025 |
| *sgs1Δ* | *sgs1Δ::loxP* | BG14 | Calugaru et al., 2025 |
| *exo1Δ sgs1Δ* | *exo1Δ::loxP sgs1Δ::loxP* | BG14 | This work |
| *exo1Δ rad24Δ* | *exo1Δ::loxP rad24Δ::loxP* | BG14 | This work |
| *sgs1Δ rad24Δ* | *sgs1Δ::loxP rad24Δ::loxP* | BG14 | This work |
| *exo1Δ sgs1Δ rad24Δ* | *exo1Δ::loxP sgs1Δ::loxP rad24Δ::loxP* | BG14 | This work |
| *rnh1Δ* | *rnh1Δ::loxP* | BG14 | This work |
| *rnh201Δ* | *rnh201Δ::loxP* | BG14 | This work |
| *rnh1Δ rnh201Δ* | *rnh1Δ::loxP rnh201Δ::loxP* | BG14 | This work |
| *exo1Δ rnh1Δ* | *exo1Δ::loxP rnh1Δ::loxP* | BG14 | This work |
| *exo1Δ rnh201Δ* | *exo1Δ::loxP rnh201Δ::loxP* | BG14 | This work |
| *exo1Δ rnh1Δ rnh201Δ* | *exo1Δ::loxP rnh1Δ::loxP rnh201Δ::loxP* | BG14 | This work |
| *rif1Δ* | *rif1Δ::loxP* | BG14 | This work |
| *rev7Δ* | *rev7Δ::loxP* | BG14 | This work |
| *exo1Δ rev7Δ* | *exo1Δ::loxP rev7Δ::loxP* | BG14 | This work |
| *cdc13Δ** | pCU-Cdc13-$FG_3$ *cdc13Δ::loxP* [b] | BG14 | This work |
| *cdc13Δ* exo1Δ* | pCU-Cdc13-$FG_3$ *cdc13Δ::loxP exo1Δ::loxP* | BG14 | This work |
| *cdc13Δ* rad24Δ* | pCU-Cdc13-$FG_3$ *cdc13Δ::loxP rad24Δ::loxP* | BG14 | This work |
| *cdc13Δ* sgs1Δ* | pCU-Cdc13-$FG_3$ *cdc13Δ::loxP sgs1Δ::loxP* | BG14 | This work |
| *cdc13Δ* exo1Δ sgs1Δ* | pCU-Cdc13-$FG_3$ *cdc13Δ::loxP exo1Δ::loxP sgs1Δ::loxP* | BG14 | This work |
| *cdc13Δ* exo1Δ rad24Δ* | pCU-Cdc13-$FG_3$ *cdc13Δ::loxP exo1Δ::loxP rad24Δ::loxP* | BG14 | This work |
| *cdc13Δ* sgs1Δ rad24Δ* | pCU-Cdc13-$FG_3$ *cdc13Δ::loxP sgs1Δ::loxP rad24Δ::loxP* | BG14 | This work |
| *cdc13Δ* exo1Δ sgs1Δ rad24Δ* | pCU-Cdc13-$FG_3$ *cdc13Δ::loxP exo1Δ::loxP sgs1Δ::loxP rad24Δ::loxP* | BG14 | This work |
| *cdc13Δ* mec3Δ* | pCU-Cdc13-$FG_3$ *cdc13Δ::loxP mec3Δ::loxP* | BG14 | This work |
| *cdc13Δ* exo1Δ mec3Δ* | pCU-Cdc13-$FG_3$ *cdc13Δ::loxP exo1Δ::loxP mec3Δ::loxP* | BG14 | This work |
| *cdc13Δ* sgs1Δ mec3Δ* | pCU-Cdc13-$FG_3$ *cdc13Δ::loxP sgs1Δ::loxP mec3Δ::loxP* | BG14 | This work |

| | | | |
|---|---|---|---|
| *cdc13Δ* exo1Δ sgs1Δ mec3Δ* | pCU-Cdc13-$FG_3$ *cdc13Δ::loxP exo1Δ::loxP sgs1Δ::loxP mec3Δ::loxP* | BG14 | This work |
| *cdc13Δ* rad9Δ* | pCU-Cdc13-$FG_3$ *cdc13Δ::loxP rad9Δ::loxP* | BG14 | This work |
| *cdc13Δ* exo1Δ rad9Δ* | pCU-Cdc13-$FG_3$ *cdc13Δ::loxP exo1Δ::loxP rad9Δ::loxP* | BG14 | This work |
| *cdc13Δ* sgs1Δ rad9Δ* | pCU-Cdc13-$FG_3$ *cdc13Δ::loxP sgs1Δ::loxP rad9Δ::loxP* | BG14 | This work |
| *cdc13Δ* exo1Δ sgs1Δ rad9Δ* | pCU-Cdc13-$FG_3$ *cdc13Δ::loxP exo1Δ::loxP sgs1Δ::loxP rad9Δ::loxP* | BG14 | This work |
| *cdc13Δ* rnh1Δ rnh201Δ* | pCU-Cdc13-$FG_3$ *cdc13Δ::loxP rnh1Δ::loxP rnh201Δ::loxP* | BG14 | This work |
| *cdc13Δ* exo1Δ rnh1Δ rnh201Δ* | pCU-Cdc13-$FG_3$ *cdc13Δ::loxP exo1Δ::loxP rnh1Δ::loxP rnh201Δ::loxP* | BG14 | This work |
| *cdc13Δ* rad24Δ rnh1Δ rnh201Δ* | pCU-Cdc13-$FG_3$ *cdc13Δ::loxP rad24Δ::loxP rnh1Δ::loxP rnh201Δ::loxP* | BG14 | This work |
| *cdc13Δ* sgs1Δ rnh1Δ rnh201Δ* | pCU-Cdc13-$FG_3$ *cdc13Δ::loxP sgs1Δ::loxP rnh1Δ::loxP rnh201Δ::loxP* | BG14 | This work |
| *cdc13Δ* exo1Δ sgs1Δ rnh1Δ rnh201Δ* | pCU-Cdc13-$FG_3$ *cdc13Δ::loxP exo1Δ::loxP sgs1Δ::loxP rnh1Δ::loxP rnh201Δ::loxP* | BG14 | This work |
| *cdc13Δ* exo1Δ rad24Δ rnh1Δ rnh201Δ* | pCU-Cdc13-$FG_3$ *cdc13Δ::loxP exo1Δ::loxP rad24Δ::loxP rnh1Δ::loxP rnh201Δ::loxP* | BG14 | This work |
| *cdc13Δ* sgs1Δ rad24Δ rnh1Δ rnh201Δ* | pCU-Cdc13-$FG_3$ *cdc13Δ::loxP sgs1Δ::loxP rad24Δ::loxP rnh1Δ::loxP rnh201Δ::loxP* | BG14 | This work |
| *cdc13Δ* exo1Δ sgs1Δ rad24Δ rnh1Δ rnh201Δ* | pCU-Cdc13-$FG_3$ *cdc13Δ::loxP exo1Δ::loxP sgs1Δ::loxP rad24Δ::loxP rnh1Δ::loxP rnh201Δ::loxP* | BG14 | This work |
| *cdc13Δ* rad24Δ rif1Δ* | pCU-Cdc13-$FG_3$ *cdc13Δ::loxP rad24Δ::loxP rif1Δ::loxP* | BG14 | This work |
| *cdc13Δ* exo1Δ rad24Δ rif1Δ* | pCU-Cdc13-$FG_3$ *cdc13Δ::loxP exo1Δ::loxP rad24Δ::loxP rif1Δ::loxP* | BG14 | This work |
| *cdc13Δ* sgs1Δ rad24Δ rif1Δ* | pCU-Cdc13-$FG_3$ *cdc13Δ::loxP sgs1Δ::loxP rad24Δ::loxP rif1Δ::loxP* | BG14 | This work |
| *cdc13Δ* exo1Δ sgs1Δ rad24Δ rif1Δ* | pCU-Cdc13-$FG_3$ *cdc13Δ::loxP exo1Δ::loxP sgs1Δ::loxP rad24Δ::loxP rif1Δ::loxP* | BG14 | This work |
| *cdc13Δ* rev7Δ* | pCU-Cdc13-$FG_3$ *cdc13Δ::loxP rev7Δ::loxP* | BG14 | This work |
| *cdc13Δ* rad24Δ rev7Δ* | pCU-Cdc13-$FG_3$ *cdc13Δ::loxP rad24Δ::loxP rev7Δ::loxP* | BG14 | This work |
| $CDC13^{RI}$* | pCU-Cdc13-$FG_3$ *cdc13Δ::CDC13-*$FG_3$ | BG14 | This work |
| $cdc13^{R411E}$* | pCU-Cdc13-$FG_3$ *cdc13Δ::*$cdc13^{R411E}$-$FG_3$ | BG14 | This work |

| | | | |
|---|---|---|---|
| $cdc13^{R531E}$* | pCU-Cdc13-FG$_3$ cdc13Δ::$cdc13^{R531E}$-FG$_3$ | BG14 | This work |
| $cdc13^{RREE}$* | pCU-Cdc13-FG$_3$ cdc13Δ::$cdc13^{RREE}$-FG$_3$ | BG14 | This work |
| $CDC13^{RI}$* exo1Δ sgs1Δ rad24Δ | pCU-Cdc13-FG$_3$ cdc13Δ::CDC13-FG$_3$ exo1Δ sgs1Δ rad24Δ | BG14 | This work |
| $cdc13^{R411E}$* exo1Δ sgs1Δ rad24Δ | pCU-Cdc13-FG$_3$ cdc13Δ::$cdc13^{R411E}$-FG$_3$ exo1Δ sgs1Δ rad24Δ | BG14 | This work |
| $cdc13^{R531E}$* exo1Δ sgs1Δ rad24Δ | pCU-Cdc13-FG$_3$ cdc13Δ::$cdc13^{R531E}$-FG$_3$ exo1Δ sgs1Δ rad24Δ | BG14 | This work |
| $cdc13^{RREE}$* exo1Δ sgs1Δ rad24Δ | pCU-Cdc13-FG$_3$ cdc13Δ::$cdc13^{RREE}$-FG$_3$ exo1Δ sgs1Δ rad24Δ | BG14 | This work |

[a] All the knockout strains were generated using a disruption cassette containing the recyclable $NAT^R$ marker from pBV65, flanked by the appropriate upstream and downstream sequences of the genes to be disrupted. Recycling of the $NAT^R$ marker by Cre-mediated recombination results in loss of the marker, leaving a LoxP site in place of the ORF.

[b] The conditional *cdc13Δ** strain was generated by first introducing the pCU-Cdc13-FG$_3$ plasmid (carrying a tagged, wild type *CDC13* gene) into the BG14 strain, followed by disruption of the chromosomal *CDC13* allele.

**Supplementary Table 2. Oligonucleotides used in this study**

| Name | Sequence 5' to 3' |
|---|---|
| ***Expression vectors*** | |
| CgCdc13-aa1-F-Bam | TTA GGATCC ATG tct att gag atc ata gcc gtg |
| CgCdc13-aa753-R-Not | AA GCGGCCGC ttg agt taa cga caa ggt ttc |
| CgCdc13-aa1-F-XmaI | TTCCCGGGATGTCTATTGAGATCATAGCCGTG |
| SMT3-FG-dn-xho | gtgctcgaGTTACTTGTCATCG |
| Gibson-Cdc13FG-pCN-F | GGATGACGATGACAAGTAA ctcgag aacacaacccacag |
| Gibson-Cdc13FG-pCN-R | GCTATGATCTCAATAGACAT cccggg ggatccactagttc |
| ***Gene disruption and mutagenesis*** | |
| Cdc13-5UTR-F3-Hpa | TTCGTTAACCGTCTACTCTGTAGAGTCTAGAATG |
| Cdc13-5UTR-R2-SacII | TTACCGCGGcacaCCAAAAGAATATCAGAACCG |
| Cdc13-3UTR-F-PvuII | TTACAGCTGGCTTTTGCTTAGCATCGAAGTG |
| Cdc13-3UTR-R-Aat | GTAGACGTCACACACAAGACTTC |
| Mec3-Aat-5UTR-F | cccgaaaagtgccacctgacgtc CCTAGCAAGTGCACATACTG |
| Mec3-Aat-5UTR-R | CAGTATGTGCACTTGCTAGG gacgtcaggtggcacttttcggg |
| Mec3-Pvu-5UTR-R | cctgcagcgtacgaagcttcagctg CCTGCCGTTACAATTGGC |
| Mec3-Pvu-5UTR-F | GCCAATTGTAACGGCAGG cagctgaagcttcgtacgctgcagg |
| Mec3-SacII-3UTR-F | ctagtggcctatgcggccgcgg ATGCATAGCTGTTTGGACTT |
| Mec3-SacII-3UTR-R | AAGTCCAAACAGCTATGCAT ccgcggccgcataggccactag |
| Mec3-HpaI-3UTR-R | ggccgattcattaatgcaggttaac TTACCTTGCTCACGAGTG |
| Mec3-HpaI-3UTR-F | CACTCGTGAGCAAGGTAA gttaacctgcattaatgaatcggcc |
| Mre11-Aat-5UTR-F | cccgaaaagtgccacctgacgtc CTTCAGCAAGGTTGCTG |
| Mre11-Aat-5UTR-R | CAGCAACCTTGCTGAAG gacgtcaggtggcacttttcggg |
| Mre11-Pvu-5UTR-R | cctgcagcgtacgaagcttcagctg TTGTCGTTGTATGCTGTTTG |
| Mre11-Pvu-5UTR-F | CAAACAGCATACAACGACAA cagctgaagcttcgtacgctgcagg |
| Mre11-SacII-3UTR-F | ctagtggcctatgcggccgcgg CTTTGGAATTACTATGCATACG |
| Mre11-SacII-3UTR-R | CGTATGCATAGTAATTCCAAAG ccgcggccgcataggccactag |
| Mre11-EcoRI-3UTR-R | ggccgattcattaatgcaggaattc GTGTGGTTCATTCATCATTCC |
| Mre11-EcoRI-3UTR-F | GGAATGATGAATGAACCACAC gaattcctgcattaatgaatcggcc |
| Sgs1-Aat-5UTR-F | cccgaaaagtgccacctgacgtc TACTACGTGGAAAAGACCG |
| Sgs1-Aat-5UTR-R | CGGTCTTTTCCACGTAGTA gacgtcaggtggcacttttcggg |
| Sgs1-Pvu-5UTR-R | cctgcagcgtacgaagcttcagctg ATCACTTTGGTGGATTATTAATTG |
| Sgs1-Pvu-5UTR-F | CAATTAATAATCCACCAAAGTGAT cagctgaagcttcgtacgctgcagg |
| Sgs1-SacII-3UTR-F | ctagtggcctatgcggccgcgg GCGCATATAAATAGAAAAAGTTAAAG |
| Sgs1-SacII-3UTR-R | CTTTAACTTTTTCTATTTATATGCGC ccgcggccgcataggccactag |
| Sgs1-HpaI-3UTR-R | ggccgattcattaatgcaggttaac GTGAAGCATTGTGCTCAC |
| Sgs1-HpaI-3UTR-F | GTGAGCACAATGCTTCAC gttaacctgcattaatgaatcggcc |

| | |
|---|---|
| Rad9_AatII-5UTR-F | aagtgccacctgacgtc CCATTCAATGCATAGAATTGTCG |
| Rad9_AatII-5UTR-R | CGACAATTCTATGCATTGAATGG gacgtcaggtggcactt |
| Rad9_PvuII-5UTR-R | gcgtacgaagcttcagctg ACCACTCTAGATTGAAAATTTCTAC |
| Rad9_PvuII-5UTR-F | GTAGAAATTTTCAATCTAGAGTGGT cagctgaagcttcgtacgc |
| Rad9_SacII-3UTR-F | ctatgcggccgcgg AGCTCTTAAACTCTGATCTTTTG |
| Rad9_SacII-3UTR-R | CAAAAGATCAGAGTTTAAGAGCT ccgcggccgcatag |
| Rad9_HpaI-3UTR-R | ccgattcattaatgcaggttaac CGAATCGAGACGATGGTC |
| Rad9_HpaI-3UTR-F | GACCATCGTCTCGATTCG gttaacctgcattaatgaatcgg |
| Rif1_AatII-5UTR-F | aagtgccacctgacgtc CCGAGTACCCCTCCATA |
| Rif1_AatII-5UTR-R | TATGGAGGGGTACTCGG gacgtcaggtggcactt |
| Rif1_PvuII-5UTR-R | gcgtacgaagcttcagctg GGTACCTATATCACTGATGTTGC |
| Rif1_PvuII-5UTR-F | GCAACATCAGTGATATAGGTACC cagctgaagcttcgtacgc |
| Rif1_SacII-3UTR-F | ctatgcggccgcgg CCACTTACTGTTAAGATTTGAAACC |
| Rif1_SacII-3UTR-R | GGTTTCAAATCTTAACAGTAAGTGG ccgcggccgcatag |
| Rif1_HpaI-3UTR-R | ccgattcattaatgcaggttaac CAACACCAGAATCAGAGCA |
| Rif1_HpaI-3UTR-F | TGCTCTGATTCTGGTGTTG gttaacctgcattaatgaatcgg |
| CgRev7-Aat-5UTR-F | cccgaaaagtgccacct gacgtc CACCAATGGATTATACAATAAG |
| CgRev7-Pvu-5UTR-R | cctgcagcgtacgaagctt cagctg TCATTTAACTTCCAAGTAAAG |
| CgRev7-Pvu-5UTR-F | CTTTACTTGGAAGTTAAATGA cagctg aagcttcgtacgctgcagg |
| CgRev7-SacII-3UTR-R | TCTTACAAGCTCATAATGGC ccgcgg ccgcataggccactagtgg |
| CgRev7-SacII-3UTR-F | ctagtggcctatgcgg ccgcgg GCCATTATGAGCTTGTAAGATATG |
| CgRev7-HpaI-3UTR-R | ccgattcattaatgcag gttaac ACACCGGGTACTGTGTCAAGC |
| CgRev7-HpaI-3UTR-F | CTTGACACAGTACCCGGTGTgttaac ctgcattaatgaatcggcc |
| CgRev7-Aat-5UTR-R | CTTATTGTATAATCCATTGGTG gacgtc aggtggcacttttcggg |
| CgRNH1-Aat-5UTR-F | cccgaaaagtgccacct gacgtc CAGCTACTTATCCATAATCGTTG |
| CgRNH1-Pvu-5UTR-R | cctgcagcgtacgaagctt cagctg TCTTGTCAAGTTCTTTGTGAT |
| CgRNH1-Pvu-5UTR-F | ATCACAAAGAACTTGACAAGA cagctg aagcttcgtacgctgcagg |
| CgRNH1-SacII-3UTR-R | TACGCACATAGCCTACCAGA ccgcgg ccgcataggccactagtgg |
| CgRNH1-SacII-3UTR-F | ctagtggcctatgcgg ccgcgg TCTGGTAGGCTATGTGCGTATATTC |
| CgRNH1-HpaI-3UTR-R | ccgattcattaatgcag gttaac AACCTCATATGTTGAAGAATGCTTTC |
| CgRNH1-HpaI-3UTR-F | GCATTCTTCAACATATGAGGTT gttaac ctgcattaatgaatcggcc |
| CgRNH1-Aat-5UTR-R | CAACGATTATGGATAAGTAGCTG gacgtc aggtggcacttttcggg |
| Rnh2_AatII-5UTR-F | aagtgccacct gacgtc CCCAGCGGTGATATCTC |
| Rnh2_AatII-5UTR-R | GAGATATCACCGCTGGG gacgtc aggtggcactt |
| Rnh2_PvuII-5UTR-R | gcgtacgaagctt cagctg ACTTAATCGATCTATAAAGTATCTTGAC |
| Rnh2_PvuII-5UTR-F | GTCAAGATACTTTATAGATCGATTAAGT cagctg aagcttcgtacgc |
| Rnh2_SacII-3UTR-F | ctatgcgg ccgcgg AGCTCCAGATGGCAAG |
| Rnh2_SacII-3UTR-R | CTTGCCATCTGGAGCT ccgcgg ccgcatag |
| Rnh2_HpaI-3UTR-R | ccgattcattaatgcag gttaac CCACACAGCCAAGTTAC |
| Rnh2_HpaI-3UTR-F | GTAACTTGGCTGTGTGG gttaac ctgcattaatgaatcgg |

| | |
|---|---|
| Cdc13-R531E-F | TATGGAATACTTTGTGAAATAGTGATGAAGTGTAAACTATATCAATCT |
| Cdc13-R531E-R | ACACTTCATCACTATTTCACAAAGTATTCCATACTTACTAACATTTTT |
| Cdc13-R411E-F | ATCGATGAGTCTACTGAGGATCAAGTCATTCAATTTAAGGACATGATA |
| Cdc13-R411E-R | TTGAATGACTTGATCCTCAGTAGACTCATCGATGAAATT TATACTGTT |
| **Vectors for FLAG3 tagging** | |
| pBV_CgCdc13-C_F | catatgggcggccgcaACGGCGATATAGCATCCTCG |
| FG_CgCdc13-C_R | GTAGTCtgccgctgcTTGAGTTAACGACAAGGTTTC |
| CgCdc13-C_FG_F | CGTTAACTCAAgcagcggcaGACTACAAAGACCATGACGG |
| CgCdc13-3UTR_pBV_R | GATGCTAAGCAAAAGCataggccactagtggatctg |
| pBV_CgCdc13-3UTR_F | gatccactagtggcctatGCTTTTGCTTAGCATCGAAGTG |
| pBV_CgCdc13-3UTR_R | gcagatccgcggccgcAGTAGAGAGCAAAAgggaaaaaat |
| CgCdc13-3UTR_pBV_F | ttcccTTTTGCTCTCTACTgcggccgcggatctgccggtc |
| CgCdc13-C_pBV_R | GATGCTATATCGCCGTtgcggccgcccatatggtgc |
| **Hybridization and Telomere PCR** | |
| CgTEL-G3 | [TGTGGGGTCTGGGTGC]$_3$ |
| CgTEL-C3 | [GCACCCAGACCCCACA]$_3$ |
| CgC3-probe-tail | [GCACCCAGACCCCACA]$_3$ TAGATAGTTGAGAGTC |
| CgG3-probe-tail | AAATAA [TGTGGGGTCTGGGTGC]$_3$ AATAA TAGATAGTTGAGAGTC |
| Cg-non-Rad-primer | GACTCTCAACTATCTA |
| CgC-telorette-1 | GCTCCGTGCATCTGGCATC CCCACAG |
| CgC-telorette-2 | GCTCCGTGCATCTGGCATC CCACAGC |
| CgC-telorette-3 | GCTCCGTGCATCTGGCATC CACAGCA |
| CgC-telorette-4 | GCTCCGTGCATCTGGCATC ACAGCAC |
| CgC-telorette-5 | GCTCCGTGCATCTGGCATC CAGCACC |
| CgC-telorette-6 | GCTCCGTGCATCTGGCATC AGCACCC |
| CgC-telorette-7 | GCTCCGTGCATCTGGCATC GCACCCA |
| CgC-telorette-8 | GCTCCGTGCATCTGGCATC CACCCAG |
| CgC-telorette-9 | GCTCCGTGCATCTGGCATC ACCCAGA |
| CgC-telorette-10 | GCTCCGTGCATCTGGCATC CCCAGAC |
| CgC-telorette-11 | GCTCCGTGCATCTGGCATC CCAGACC |
| CgC-telorette-12 | GCTCCGTGCATCTGGCATC CAGACCC |
| CgC-telorette-13 | GCTCCGTGCATCTGGCATC AGACCCC |
| CgC-telorette-14 | GCTCCGTGCATCTGGCATC GACCCCA |
| CgC-telorette-15 | GCTCCGTGCATCTGGCATC ACCCCAC |
| CgC-telorette-16 | GCTCCGTGCATCTGGCATC CCCCACA |
| CgSubtel-F1 | GGGTTAAATAGGGTTGTGTAGGGTTCC |
| CgSubtel-F4 | CCCATATATGCCTTGATCACTTGTCACC |
